# Distribution and molecular evolution of the anti-CRISPR family AcrIF7

**DOI:** 10.1101/2021.06.27.450086

**Authors:** Wendy Figueroa, Adrián Cazares, Daniel Cazares, Yi Wu, Ana de la Cruz, Martin Welch, Luis Kameyama, Franklin L. Nobrega, Gabriel Guarneros

**Author notes:** These authors contributed equally (Wendy Figueroa, Adrián Cazares).

## Abstract

Anti-CRISPRs are proteins capable of blocking CRISPR-Cas systems and typically encoded in mobile genetic elements. Since their discovery, numerous anti-CRISPR families have been identified. However, little is known about the distribution and sequence diversity of members within a family, nor how these traits influence the anti-CRISPR’s function and evolution. Here we use AcrIF7 to explore the dissemination and molecular evolution of an anti-CRISPR family. We uncovered five sub-clusters and prevalent anti-CRISPR variants within the group. Remarkably, AcrIF7 homologs display high similarity despite their broad geographical, ecological and temporal distribution. Although mainly associated with *Pseudomonas aeruginosa*, AcrIF7 was identified in distinct genetic backgrounds indicating horizontal dissemination, primarily by phages. Using mutagenesis, we recreated variation observed in databases but also extended the sequence diversity of the group. Characterisation of the variants identified residues key for the anti-CRISPR function and other contributing to its mutational tolerance. Moreover, molecular docking revealed that variants with affected function lose key interactions with its CRISPR-Cas target. Analysis of publicly available data and the generated variants suggests that the dominant AcrIF7 variant corresponds to the minimal and optimal anti-CRISPR selected in the family. Our study provides a blueprint to investigate the molecular evolution of anti-CRISPR families.

## INTRODUCTION

Bacteria are constantly under attack by bacteriophages. As a result, they have evolved an extensive array of anti-phage defence systems such as abortive infection, restriction-modification, and CRISPR-Cas (an acronym for Clustered Regularly Interspaced Short Palindromic Repeats-CRISPR-associated proteins) [1]. CRISPR-Cas systems are a type of prokaryotic adaptive immune system in which specific sequences targeting foreign nucleic acids are integrated into the bacterial chromosome and provide protection against exogenic mobile elements [2–5]. These systems, widespread amongst bacteria [6], detect the invader nucleic acid prior its degradation via an endonuclease. In the opportunistic pathogen *Pseudomonas aeruginosa*, three different subtypes of the CRISPR-Cas system have been described to date: I-C, I-E and I-F, with the latter being the most abundant [7, 8].

In response to the infection barriers deployed by their hosts, phages have developed various mechanisms to evade defence systems, leading to a dynamic evolutionary arms race [9]. One strategy that phages evolved to circumvent CRISPR-Cas immunity is the use of proteins that block the system, known as anti-CRISPRs (Acr). These proteins, first described in *P. aeruginosa* phages evading the type I-F and I-E systems [10, 11], have been identified in mobile genetic elements such as phages and plasmids, but are also encoded in bacterial genomes. In fact, it has been proposed that Acr proteins are present in more than 30% of *P. aeruginosa* strains that carry CRISPR-Cas systems [8].

Since their discovery, anti-CRISPR research has mostly centred on identifying new anti-CRISPR genes in diverse bacterial species [12, 13]. Around 90 anti-CRISPR families evading different CRISPR-Cas system types have been reported [14] (http://cefg.uestc.cn/anti-CRISPRdb/). These families rarely share sequence similarity and seem to possess distinctive molecular mechanisms of action. As the number of reported anti-CRISPRs keeps rising, efforts have been put into compiling and organising anti-CRISPR sequences and their metadata in the form of a resource database [12]. In its first version, anti-CRISPRdb contained 432 entries, including both experimentally validated and computationally predicted anti-CRISPRs. Currently, this database holds information for more than 3,600 anti-CRISPR proteins corresponding to at least 85 Acr sub-types (http://cefg.uestc.cn/anti-CRISPRdb/).

In comparison to the comprehensive efforts put into discovering anti-CRISPR families, the mechanisms of action of these powerful molecules have been investigated to a much lesser extent. Most of the Acr mechanisms characterised to date involve blockage of different steps in the CRISPR-Cas restriction process: DNA binding, cleavage, crRNA loading, or formation of the effector-complex [15–17]. For example, AcrIF1 binds Csy3 blocking the recognition of the invading DNA [15], whereas AcrIF2 and AcrIF7 interact with the Csy1-Csy2 complex impeding DNA binding [15, 18]. Additionally, a few anti-CRISPR structural studies have described inter-protein interactions occurring with different components of the CRISPR-Cas system. Some of these reports addressed the identification of important residues in the protein, typically by changing amino acids such as alanine from polar to non-polar by site-directed mutagenesis [18, 19].

Besides their molecular mechanism, the evolution of anti-CRISPR families has remained largely unexplored so far. Little is known about the distribution and sequence diversity of anti-CRISPRs belonging to the same family. Still, these attributes are key to understanding how anti-CRISPRs of a certain type are acquired, what their host range is, to what extent their sequences have changed and whether such changes impact the protein function.

Here we use AcrIF7 as a model to study the molecular evolution of an anti-CRISPR family. We uncovered the AcrIF7 diversity and distribution by analysing homologs identified in bacterial and phage genomes. We report prevalent sequence variants and show that AcrIF7 homologs display high similarity despite their occurrence in diverse genome regions and wide geographical, ecological and temporal distribution. Using random and site-directed mutagenesis, we generated observed and novel AcrIF7 variants to investigate the impact of sequence variation in the anti-CRISPR function. Our experimental and computational characterisation discovered key residues for the anti-CRISPR function but also distinguished regions contributing to the mutational robustness of the protein. Together, our findings suggest that the dominant AcrIF7 variant represents both the optimal and minimal functional unit of the group and reveal features of AcrIF7 that can be used in favour of its development for biotechnology applications. Furthermore, our study serves as a blueprint to investigate the molecular evolution of other anti-CRISPR families.

## METHODS

### Bacterial strains, phages, and culture conditions

PA14 wild-type strain, a mutant lacking the CRISPR-Cas system (PA14 ΔCRISPR loci *Δcas* genes, also referred to as PA14ΔCR), and the phage JBD18 were kindly provided by Professor Alan R. Davidson [20]. Phage H70 harbouring *acrIF7* gene (named *g2* in the annotation of the phage genome) was isolated from a clinical *P. aeruginosa* strain [21] and belonged to Dr Gabriel Guarneros’ phage collection, along with phages Ps45 and H68. Overnight cultures were grown routinely in LB (Lennox) broth with shaking at 37°C unless otherwise indicated.

### Phage propagation and purification

Phages H70 (G2 carrier) and JBD18 (CRISPR-sensitive) were propagated and purified following the protocol previously reported by Cazares *et al*. [21]. Briefly, the phages were propagated using the standard soft agar overlay method, followed by concentration with PEG and purification by CsCl gradient centrifugation.

### Analysis of AcrIF7 sequences

The amino acid sequence of the anti-CRISPR G2 from phage H70 was first compared to all sequences available in anti-CRISPRdb [12] in August 2020 using BLASTp [22]. Only proteins of the AcrIF7 family matched G2. The 68 sequences of proteins of the AcrIF7 family available in the database were clustered using CD-HIT with an identity threshold of 100%, word size of 5 and length difference cutoff of 0 to remove identical sequences. The search of sequences homologous to G2 was extended to 50,457 proteins encoded by 574 *Pseudomonas* phage genomes available in GenBank, and 32,262,482 proteins from 5,279 *P. aeruginosa* genomes deposited in the GenBank RefSeq database in August 2020. A maximum e-value of 1e-03 was considered to identify homologs in all the BLASTp searches.

The amino acid sequences of G2, the 119 homologs identified in *Pseudomonas* phages and *P. aeruginosa* genomes, and the 25 non-redundant AcrIF7 sequences retrieved from anti-CRISPRdb, were aligned using the PRALINE algorithm [23] with default settings. A neighbour-joining tree was inferred from the multiple sequence alignment using the BioNJ method integrated into Seaview v4.6 [24] with 1,000 bootstrap replicates, observed distance, and including gap sites. The resulting tree was visualised with iTOL v5.7[25]. An alignment of non-redundant sequences selected from the tree (deduplicated with CD-HIT and aligned with PRALINE as described above) was visualised and edited with Jalview v2.11.1.3 [26]. Protein sequences representative of each sub-cluster identified in the neighbour-joining tree were scanned against all member databases in InterPro using InterProScan v5.50-84.0 [27] with default settings. The search for homologs in non-*P. aeruginosa* genomes was performed at the nucleotide level in the BLASTn suite online [22] with default search parameters, excluding *P. aeruginosa* (taxid:287) in the organisms list, and using a representative of each AcrIF7 sub-cluster as query. AcrIF7 homologs in plasmids were searched with BLASTn against the pATLAS database [28].

Nucleotide sequences of *g2* homologs from the sub-clusters sc1 and sc2 identified in the tree were extracted from the corresponding genomic fasta files and aligned at the protein level with Seaview v4.6 [24] using the PRALINE alignment as a template. The resulting multiple sequence alignment at the nucleotide level was used in the positive selection analysis (see below). Additionally, a sequence stretch including the *acrIF7* CDS region plus 5 kb of the upstream and downstream sequence was extracted from *P. aeruginosa* genomes in the “complete” and “chromosome” GenBank assembly categories and phage genomes identified as carrying a G2 homolog. The extracted sequences were compared with BLASTn [22] and the pairwise comparisons visualised with the genoPlotR package v0.8.11 [29].

### Metadata, Multilocus sequence typing (MLST) and CRISPR-Cas identification

The MLST profiles of 117 *P. aeruginosa* bacterial genomes carrying AcrIF7 were identified from the pubMLST *P. aeruginosa* scheme (http://pubmlst.org/paeruginosa/) [30] using the mlst tool v.2.8 (https://github.com/tseemann/mlst) (Supplementary Table S1). BioSample records of the genomes were retrieved from GenBank using the NCBI’s Edirect [31] to extract information on the isolation source, year and country of isolation of the bacterial strains (Supplementary Table S1). The genome sequences were also analysed with cctyper v1.4.4 [32] for the identification and subtyping of CRISPR-Cas genes and arrays.

### Positive selection analysis

Positive selection analysis was performed on 131 AcrIF7 variants, including G2, the previously reported AcrIF7 (accession number ACD38920.1), and the rest of AcrIF7 variants from sub-clusters sc1 and sc2, using codeml from the PAML package v4.9 [33]. The nucleotide alignment was trimmed in the 5’ end to the length of G2 to have an alignment of the core codons (Supplementary data - AcrIF7 alignment). The tree used for the analysis was obtained from IQTree v1.3.11 [34], with the model K2+G4. We fit the data to the site models M1a (NearlyNeutral) and M2a (PositiveSelection). We then performed a likelihood ratio χ2 test of both models and determined the p-value. Finally, Empirical Bayes (EB) [33] was used to calculate the posterior probabilities for site classes and identify dN/dS values for each codon.

### Experimental Evolution of phage H70

Evolution experiments were performed by mixing an ancestral (initial) H70 phage stock and overnight cultures of either PA14 WT or PA14 ΔCR in 30 mL of fresh LB to a final concentration of 10^6^ PFU/mL and 10^6^ CFU/mL. Cultures were incubated for 16 h at 37°C with shaking and then centrifuged at 4,000 rpm for 12 min. Supernatants containing the evolved phage were filter-sterilised and titred on both PA14 and PA14ΔCR, irrespective of where they were propagated. The consequent passages (10 in total, 3 lineages each) were done by mixing the evolved phage stock with fresh overnight cultures of the same strain used in the first passage (PA14 or PA14ΔCR) to a final concentration of 10^6^ PFU/mL and 10^6^ CFU/mL followed by the same steps as in passage 1. Each titration assay was performed in triplicate and the efficiency of plating (EOP) were calculated using PA14 ΔCR as reference (titre on PA14/ titre on PA14ΔCR). P-values were calculated using a one-way ANOVA test on each dataset. DNA from samples from passage 10 was extracted and combined in one sample per strain (H70 evolved in PA14 WT and H70 evolved in PA14 ΔCR) for Illumina sequencing. Reads were mapped against the parental strain using BWA mem v0.7.17-r1188 [35] and variants were identified using pilon v1.22 with the default settings for variant calling. The percentage of each nucleotide in each position of the H70 genome was calculated based on the number of reads with that particular nucleotide and the total number of reads covering that position. The data were filtered to exclude the variants with the reference nucleotide and those representing less than 1% of the population (Supplementary Table S2).

### Random mutagenesis

Error-prone PCR [36] was used to introduce mutations in the sequence. Three different conditions were used (Supplementary Table S3), which differ in the concentration of MgCl2, MnCl2, and the number of extension cycles. PCR products were run in a 1% agarose gel to confirm the amplification of *g2* under the mutagenic conditions, purified using Sap-Exo kit, Jena Bioscience, and cloned into a modified version of pUCP24 plasmid (Supplementary Figure S1). Chemically competent DH5α *E. coli* cells were prepared and transformed following the protocol previously reported by Green & Rogers [37]. Around 900 *E. coli* colonies were picked and grown overnight in LB-Gm (15 μg/ml). Cultures were mixed in pools of 10 candidates and plasmids were extracted using Wizard® Plus SV Minipreps DNA Purification System, Promega, to have pools of plasmids with diverse variants of *g2*, although some empty vectors were also present in the mix. Pools were then electroporated into *P. aeruginosa* PA14 following the protocol described by Choi et al. [38].

### Selection of *P. aeruginosa* candidates carrying *g2* by colony blot

We established a colony blot protocol for detecting the presence of genes in *P. aeruginosa* in scale using a radioactive probe (Supplementary Figure S3). *P. aeruginosa* colonies carrying *g2* (not empty vectors) were selected by colony blot to make the screening more efficient than by PCR. A hundred candidates were streaked on two LB-Gm (50 μg/ml) plates (a master plate and a replica plate), along with negative and positive controls (colony with empty plasmid and with *g2*, respectively), and incubated overnight at 37°C. Colonies were transferred from the replica plate to a nylon membrane. Membranes were placed onto filter paper damped with solution I (0.5M NaOH, 1.5M NaCl) for 10 min, and then placed onto another filter paper damped with solution II (1M Tris-HCl pH 7.2) for 2 min to neutralise the reaction.

Finally, the membranes were placed on the top of filter paper moistened with solution III (0.5M Tris-HCl, 1.5M NaCl) for 5 min, and exposed to UV light for 5 min in a crosslinker to fix the DNA to the membrane. The membranes were then introduced in hybridisation tubes with 10 ml of 1% SDS, 1M NaCl solution and incubated at 42°C for 2 hours. During this incubation period, an oligonucleotide complementary to *g2* (G2 exp forward, Supplementary Table S4) was labelled with Phosphorus-32 following the protocol reported by Novogrodsky *et al*. [39]. The probe was then added to the membranes and incubated at 50°C overnight. The membranes were subsequently washed with 10 ml of 2X SSC for 3 min at room temperature, followed by 2 washes with 2X SSC containing 1% SDS for 5 minutes each. The membranes were then washed for 30 min with 10 ml of 1X SSC solution, and finally with 10 ml of 0.5X SSC for 15 min. The membranes were allowed to dry before placing them onto an x-ray film in a film cassette. The film was developed, and dark spots on the film produced by the radioactive probe indicated the presence of *g2* in the colony (Supplementary Figure S3).

### Phage infection assay

A hundred microliters of overnight cultures of *P. aeruginosa* colonies carrying *g2* were mixed with 3.5 mL of TΦ top agar (1% peptone, 0.5% NaCl, 0.7% agar, 10mM MgSO4) and poured over TΦ plates (1% peptone, 0.5% NaCl, 1.5% agar) containing 50 μg/ml of gentamicin. Serial dilutions of JBD18 phage stock were spotted onto the lawns and the plates were incubated overnight at 37°C. The EOP was calculated as the titre of the JBD18 phage in PA14 carrying the variant of G2 divided by the titre in PA14 harbouring the wild-type version of G2. Each infection assay was performed in triplicate. Welch’s t-test was used and p-values were corrected for multiple comparisons using the Bonferroni correction (Supplementary Table S5).

### Sequencing and analysis of variants

Colony PCR of the colonies of *P. aeruginosa* carrying the variants of *g2* was performed using the oligos MCS pUCP24 forward and MCS pUCP24 reverse (Supplementary Table S4). The PCR products were cleaned using the Sap-Exo kit, Jena Bioscience, according to the manufacturer’s specifications and the variants were sequenced using the BigDye™ Terminator v1.1 Cycle Sequencing Kit, ThermoFisher Scientific (Supplementary Data - Electropherograms).

### Site-directed mutagenesis

Site-directed mutagenesis of *g2* was done using the Q5® Site-Directed Mutagenesis Kit, New England Biolabs, according to the manufacturer’s specifications, using the NEBaseChanger tool (http://nebasechanger.neb.com/) for oligo design. To generate multiple changes of amino acids in a single position, the primers were designed with one or two random nucleotides in the codon of the target residue (Supplementary Table S4 - primers V40, D29 and Y32).

### Protein modelling and docking analyses

G2 wild-type and mutants were modelled using AlphaFold2 [40]. Analysis of the models was done using open-source Pymol (Schrodinger, LLC. 2010. The PyMOL Molecular Graphics System, Version 2.4.0). The binding position of G2 or its mutants on Cas8f (PDB code 7JZX and chain A) was predicted using HADDOCK 2.4 [41], as described by Kim et al [18]. The residue-residue interactions of G2/mutants and Cas8f were analysed using Ligplot+ [42]. The secondary structure alignment of G2 and its mutants was visualised using 2dSS [43], and analysis of the models and superpositions was done using chimeraX [44].

## RESULTS

### Protein G2 of phage H70 is an anti-CRISPR of the family AcrIF7

A genomic analysis of the phage H70 isolated from the *P. aeruginosa* clinical strain HIM5 [21] revealed proto-spacers in the ORFs 14 and 28 matching regions of the CRISPR loci of *P. aeruginosa* PA14 (Figure 1A). Yet, infection assays showed that the phage H70 could infect the strain PA14 (Figure 1B). Analysis of the phage H70 accessory genome identified an anti-CRISPR locus in the region of genomic plasticity (RGP) G composed of the genes *g2* and *g9* [21], which are homologous to *acrIF7* and *aca1*, respectively [45]. AcrIF7 was first reported as an 83 aa protein (GenBank accession ACD38920.1) with anti-CRISPR activity against the CRISPR-Cas system I-F [12, 45]. In comparison, G2 of phage H70 (Accession YP_009152337.1) is 67 aa long, lacking 16 amino acids in the N-terminus of ACD38920.1.

**Figure 1.**
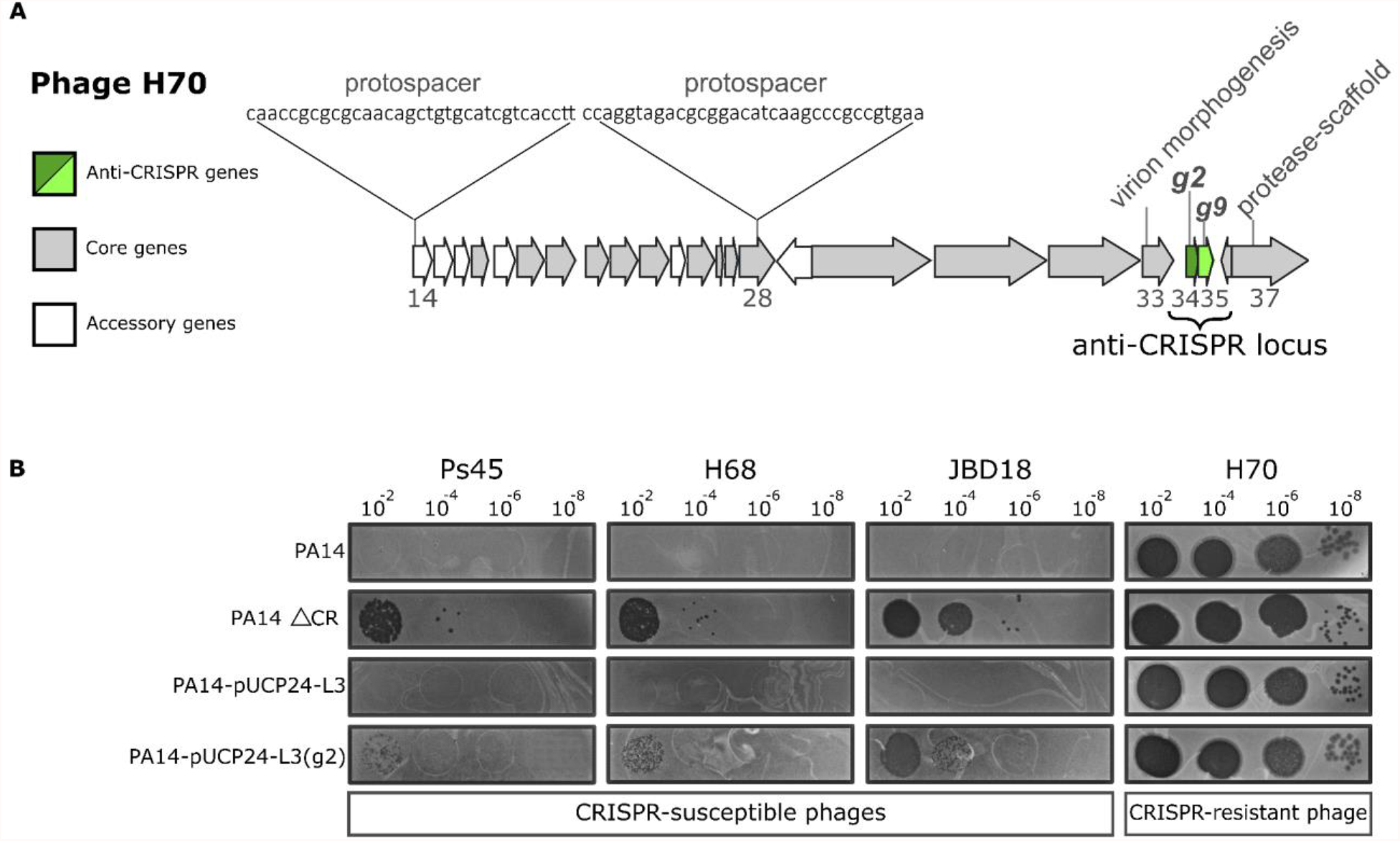
Location of the anti-CRISPR gene *g2* in the genome of phage H70 and inhibition of the CRISPR-Cas system I-F. A) The map represents a region of the H70 phage genome (ORFs 14 to 37 shown as arrows). The grey arrows correspond to core genes conserved in the phage group D3112virus, whereas the white arrows represent accessory ORFs [21]. The anti-CRISPR locus, encoding the anti-CRISPR gene (*g2*) and a putative DNA binding gene (*g9*), are shown in green. B) Serial dilutions of different CRISPR-sensitive phages (indicated above the figure) were spotted on bacterial lawns of the PA14, PA14 ΔCRISPR-cas (PA14 ΔCR), PA14-pUCP24-L3, and PA14-pUCP24-L3(*g2*) strains. Phage infection (shown as plaques) denotes a lack of CRISPR-Cas defence due to either the absence of the CRISPR-Cas system (PA14 ΔCR) or anti-CRISPR activity (PA14-pUCP24-L3(*g2*)). Note that the titre of each phage stock was different, and therefore not comparable between phages.

To test the functionality of this shorter version of AcrIF7, we cloned *g2* in the plasmid pUCP24-L3 (Supplementary Figure S1) and assessed the protection of CRISPR-sensitive phages against the CRISPR-Cas system. The infection assays were performed in the strains PA14 wild-type (WT), a PA14 mutant lacking the CRISPR loci and *cas* genes (PA14 ΔCR), PA14 carrying the plasmid with *g2* (PA14-pUCP24-L3(*g2*)), and PA14 transformed with the empty vector (PA14-pUCP24-L3). Phage H70 was able to infect all the strains with similar efficiency, regardless of the presence of the CRISPR-Cas system (Figure 1B). In contrast, CRISPR-sensitive phages Ps45, H68, and JBD18, only produced lytic plaques in the PA14 ΔCR mutant or the strain PA14 WT carrying *g2*; thus demonstrating that this shorter version of AcrIF7 is a functional anti-CRISPR (Figure 1B).

### AcrIF7 family is conserved and mainly associated with *P. aeruginosa* prophages

Since the major difference between G2 and the first reported AcrIF7 is the additional 16 amino acids in the N-terminus of ACD38920.1, we sought to investigate the diversity of this anti-CRISPR family. A comparative search against the ~3,600 sequences available in anti-CRISPRdb [12] only identified proteins of the AcrIF7 family as homologous to G2. The 68 homologs identified in the database, corresponding to 25 unique sequences, included a pair of chimeric proteins with homology to anti-CRISPR of the families IE4 and IF7. Members of the AcrIF7 family reported in anti-CRISPRdb are mostly associated with *P. aeruginosa* genomes but also found in different *Janthinobacterium* species and in *P. citronellolis*, with the latter corresponding to AcrIE4-F7 anti-CRISPR hybrids (Figure 2) [46].

**Figure 2.**
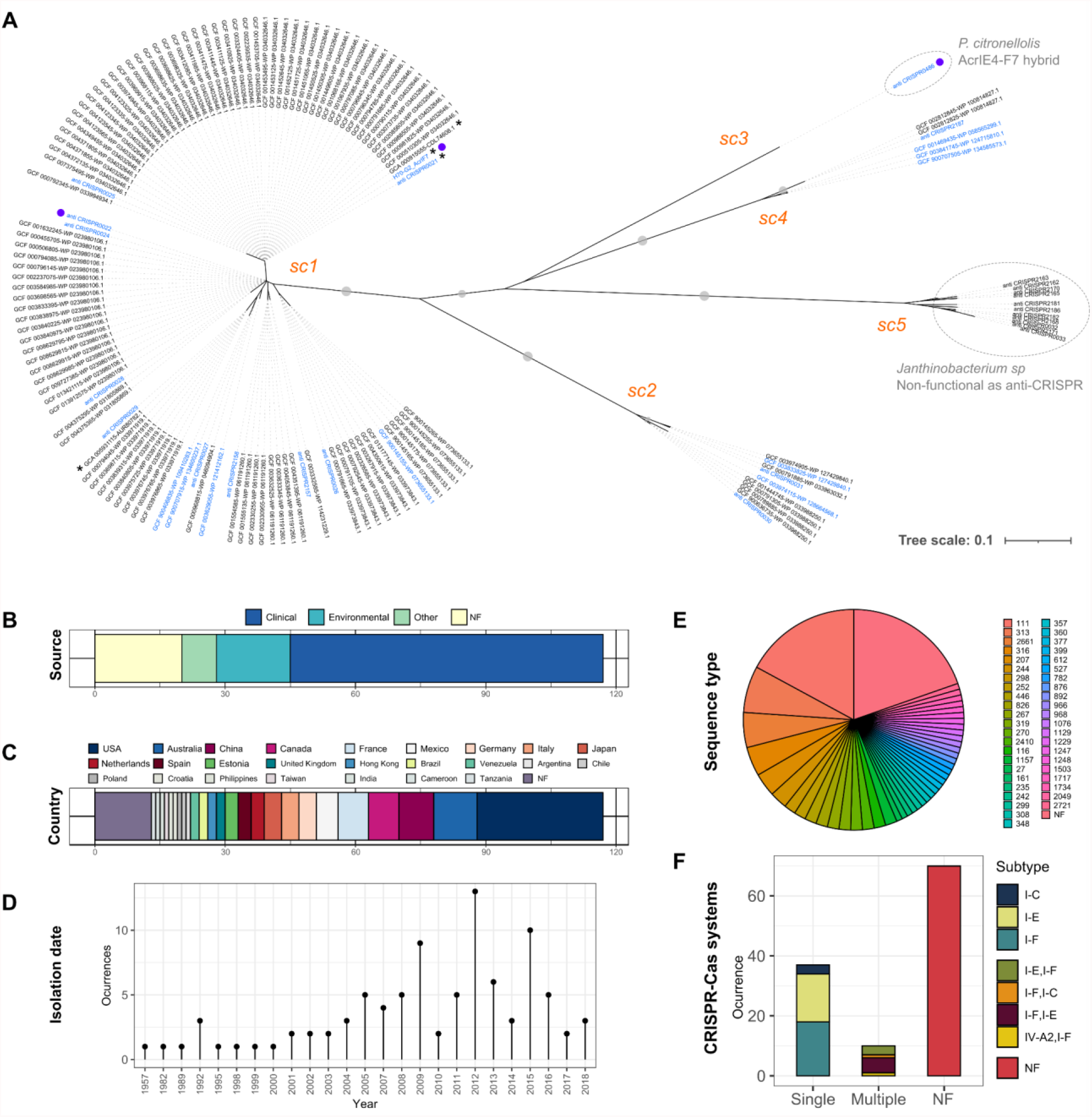
Diversity of members of the anti-CRISPR family AcrIF7. A) Neighbour-joining unrooted tree displaying the patterns of sequence similarity among protein sequences homologous to G2 of phage H70. Homologous sequences were identified through BLASTp searches against anti-CRISPRdb [12] and proteomes of *Pseudomonas* phages and *P. aeruginosa* genomes from GenBank (see Methods). The amino acid sequences of the 145 homologs presented in the tree were aligned with PRALINE [23]. The tree was inferred from the resulting alignment with Seaview v4.6 [24] (BioNJ method). Grey dots on tree branches represent bootstrap support values >80 calculated from 1000 replicates. Sub-clusters (sc) identified in the tree are indicated in orange. The 25 non-redundant sequences from anti-CRISPRdb included in the tree are labelled with their corresponding identifier in the database (“anti_CRISPR” prefix). The remaining sequence labels indicate the GenBank assembly identifier and protein accession number separated by a hyphen (“-”) except for the sequence corresponding to G2 of phage H70. Asterisks mark sequences identified in phage genomes, whereas purple dots pinpoint sequences that have been experimentally verified as an anti-CRISPR. Labels in blue denote non-redundant sequences within their corresponding sub-cluster (excluding those in sc5) and thus represent the diversity of protein sequences in the tree. Dotted line circles indicate sequences identified in non-*P. aeruginosa* genomes. Notes on the hybrid nature of the sequence in sc3, and the lack of identifiable anti-CRISPR activity against the systems IF and IE of *P. aeruginosa* in a homolog (accession: WP_034755374.1) of sc5, correspond to references [47] and [45]. B-F) Metadata associated with *P. aeruginosa* genomes encoding an AcrIF7 homolog. Data plotted in B-D were extracted from the genomes BioSample record (Supplementary Table S1). Sequence Types (ST) presented in panel E were identified from the pubMLST *P. aeruginosa* scheme [48], http://pubmlst.org/paeruginosa) using the mlst tool v.2.8 [49], https://github.com/tseemann/mlst). The occurrence of CRISPR-Cas systems in the AcrIF7-carrier genomes, displayed in panel F, was assessed with cctyper v1.4.4 [32].

To further explore the diversity and distribution of the AcrIF7 family, we expanded our homology search to all proteins encoded in *Pseudomonas* phage genomes deposited in GenBank and *P. aeruginosa* genomes available in RefSeq. One hundred nineteen homologs were identified, predominantly in *P. aeruginosa* genomes. Multilocus sequence typing (MLST) analysis of the bacterial sequences distinguished 44 types in 94 genomes, with the remaining 23 sequences missing one or two alleles, thereby highlighting the diversity of *P. aeruginosa* isolates carrying an anti-CRISPR of the AcrIF7 family (Figure 2; Supplementary Table S1). Likewise, analysis of metadata retrieved for the *P. aeruginosa* genomes revealed an extensive geographical, temporal and source distribution encompassing five continents, over six decades and a large variety of clinical and environmental samples (Figure 2).

Comparison of the newly-identified AcrIF7 homologs with the non-redundant protein sequences from anti-CRISPRdb and that from phage H70 uncovered five sub-clusters within the family (Figure 2). Three sub-clusters (sc1, sc2 and sc4) were detected in *P. aeruginosa* and *Pseudomonas* phage genomes, with sc1 representing the dominant type. A BLAST search at nucleotide level against non-*P. aeruginosa* bacterial and phage genomes in GenBank using a representative of the five subclusters only identified a match of sc3 in *P. citronellolis* (accession: CP015878.1) and confirmed that sc5 is associated with *Janthinobacterium* species. No matches were detected in plasmid sequences reported in pATLAS [28].

Excluding sc5, for which no anti-CRISPR activity against *Pseudomonas* was detected in previous experimental characterisation [45], protein sequence similarity ranged from 62 to 81% between sub-clusters (Supplementary Table S6). Notably, AcrIF7 members of the same sub-cluster display limited sequence variation, with similarity values ranging from 98 to 100%, corresponding to 5-14 mutations (Figure 3). In the dominant type sc1, containing 117 members and including G2 of phage H70, only 15 different sequences and 14 mutations were distinguished (Figure 3). Sequences within this sub-cluster varied from 59 to 87 amino acids long, with 67 aa representing the predominant length (Figure 3). Remarkably, the prevalence of variants within the dominant AcrIF7 type sc1 also differed considerably; variants represented by G2 and anti_CRISPR0024, separated by a single mutation, corresponded to 43 and 17% of the members in the sub-cluster, respectively (Figure 3).

**Figure 3.**
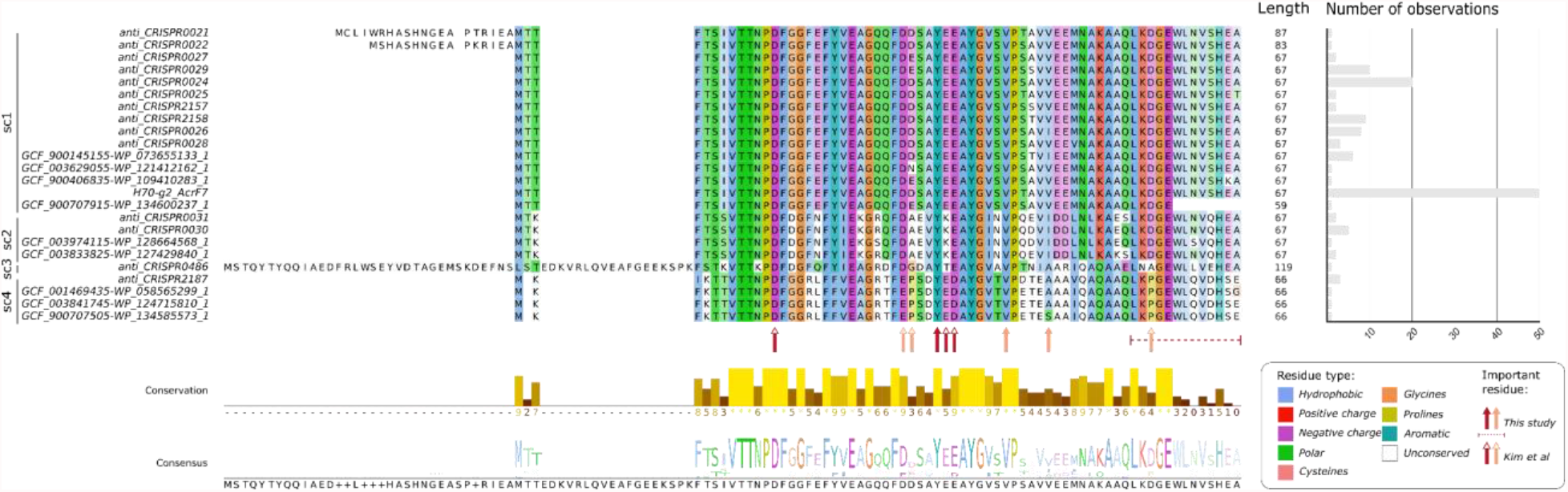
Alignment of non-redundant protein sequences of the AcrIF7 family. The 24 proteins selected as representative of the sequence diversity observed amongst *acrIF7* homologs (see sequence labels in blue in Figure 2) were aligned with PRALINE [23]. The resulting alignment was visualised with Jalview v2.11.1.4 [26]. Identifiers of the homologous variants, shown on the left side of the alignment, correspond to those described in Figure 2. The sub-cluster to which the variant belongs is indicated next to its identifier. The length of each variant sequence is displayed on the right side of the alignment, next to the bar plot illustrating the number of observations of the different variants amongst the genomes where a G2 homolog was identified (see Figure 2). Residues in the alignment are colour coded based on their level of conservation in a given position and the residue type they belong to according to the ClustalX shading scheme, indicated at the bottom-right of the figure. The conservation level and consensus sequence of the alignment are represented with a bar plot and sequence logo at the bottom of the figure, respectively. Residues identified in this study as important for the G2 anti-CRISPR activity are pinpointed with solid arrows or a dotted line below the alignment. The dotted line indicates that the lack of the underscored residues in G2 nullifies the anti-CRISPR activity of the protein (see Figure 6). Residues important for the AcrIF7-CRISPR-Cas interaction as reported by Kim et al. [18] are identified with open arrows. Red arrows denote residues on which mutations drive the loss of the AcrIF7 function or interaction whereas orange arrows indicate residues on which mutations have a partial effect.

We then investigated whether sequence conservation observed amongst homologs of the same sub-cluster was extended to the genome regions encoding the anti-CRISPR gene. We extracted and compared regions flanking anti-CRISPRs of the AcrIF7 family identified in phage and complete *P. aeruginosa* genomes. The comparison revealed that anti-CRISPRs AcrIF7 can be associated with diverse genomic backgrounds (Figure 4). Members of the sub-cluster sc1 were detected in three distinct phage types and two divergent bacterial genome regions, indicative of anti-CRISPR acquisition via horizontal gene transfer. Still, AcrIF7 members of the sub-cluster sc1 were largely linked to transposable phages of the group D3112-like (Figure 4).

**Figure 4.**
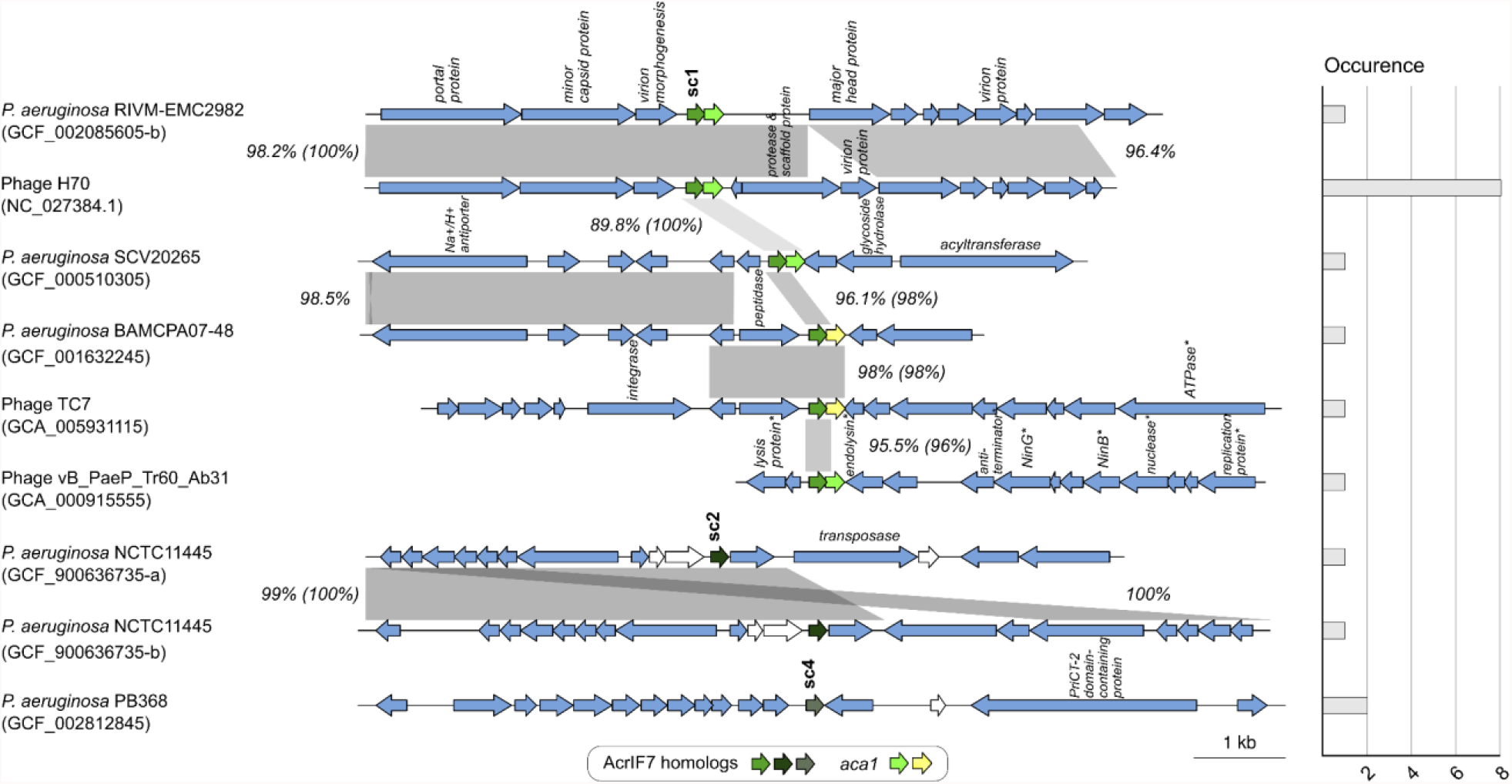
Comparative analysis of genome regions harbouring AcrIF7 homologs. The figure shows pairwise comparisons at the nucleotide level of regions containing an AcrIF7 homolog in complete *P. aeruginosa* and phage genomes. Regions containing an AcrIF7 homolog gene plus 5 kb of upstream and downstream sequence (where available), were extracted and compared all-vs-all with BLASTn. Regions were then clustered based on sequence similarity to determine their frequency of occurrence among the set of analysed genomes. We defined regions as belonging to the same type when sharing >95% overall sequence coverage and identity. The figure summarises the comparison of 17 regions from 13 genomes, displaying the similarity between representatives of the different region types identified. Regions are paired with their closest match and their occurrence is illustrated with a bar plot on the right side of the figure. The organism name and GenBank accession number (in parenthesis) of the genomes from which the regions were extracted are indicated next to their corresponding gene maps, on the left side of the figure. Instances where more than one AcrIF7 homolog was detected in the same genome, are distinguished with a suffix letter added to the GenBank accession number. The AcrIF7 homolog genes, and *aca1*, are colour coded as indicated in the figure. The sub-cluster type of the different AcrIF7 homolog genes (see Figure 2), is shown above the corresponding gene arrow. Where available, functions assigned to ORF products, as indicated in the GenBank file, are displayed above the corresponding arrow. Asterisks mark functions assigned as putative. Light yellow arrows denote ORFs encoding homologs of Aca1 overlooked in the original annotations. White arrows indicate overlooked ORFs with unknown functions. The percentage of sequence identity detected between homologous regions, depicted as grey connecting blocks, is indicated next to the corresponding block. For homologous regions containing an AcrIF7 homolog gene, the percentage of identity between the gene sequences is additionally indicated in parenthesis.

Surprisingly, the comparative analysis also uncovered the presence of more than one copy of the *acrIF7* gene in the genome of three *P. aeruginosa* strains: RIVM-EMC2982 (GCF_002085605), Carb01 63 (GCF_000981825) and NCTC11445 (GCF_900636735). In these cases, the anti-CRISPR gene was located in different genome positions, and it was frequently surrounded by homologous phage genes. In terms of neighbour genes, no other anti-CRISPRs were detected next to the *acrIF7* gene in the regions analysed, but *aca1* was typically located immediately downstream of *acrIF7* genes of the sub-cluster sc1 (Figure 4). In contrast, no homologs of Aca1, nor proteins with a lambda repressor-like DNA-binding domain characteristic of Aca1 were identified in the regions flanking members of the sub-clusters 2 and 4.

Despite the existence of sequence variation between the compared regions, the anti-CRISPR gene commonly bore no mutations or displayed nucleotide sequence conservation above the average (Figure 4). For example, the 204 bp coding region of H70 *g2* was identical in 11 genomes, including in the unrelated regions of *P. aeruginosa* SCV20265 and phage vB_Paep_Tr60_Ab31. Such levels of sequence similarity hampered the discrimination of conserved positions in the anti-CRISPR protein that could be important for its function. Screening of the AcrIF7 sequences for the presence of known protein signatures or classification into established protein families yielded no results. Thus, no functional domains or motifs, nor links to functionally characterised protein sequences were identified, underscoring the novelty of the AcrIF7 family.

### AcrIF7 family is under nearly neutral evolution

Most AcrIF7 homologs identified in GenBank displayed high protein sequence similarity (Figures 2 and 3). Yet, some positions exhibiting variation were observed when aligning the AcrIF7 non-redundant sequences (Figure 3), implying divergence at the codon level. Therefore, we sought to estimate the synonymous and nonsynonymous substitution rates of each residue, looking for codons potentially under positive selection that could drive protein fitness optimization. Only representatives of the sub-clusters sc1 and sc3 have been reported to be active anti-CRISPRs against the CRISPR-Cas system I-F of *P. aeruginosa* (Figure 2). However, since the divergence of the chimeric sc3 at the nucleotide level could compromise the reliability of the analysis, we decided to focus on the sequences of the sub-clusters sc1 and sc2. Moreover, sc1 and sc2 represent the predominant sub-clusters, accounting for around 90% of all the members in the AcrIF7 family. One hundred thirty-one sequences (Supplementary Data - AcrIF7 alignment) were analysed using the models M1a (NearlyNeutral) and M2a (PositiveSelection), which estimate the evolutionary pressures over specific codon sites in the gene (Supplementary Data - M1a/M2a). After performing a likelihood ratio χ2-test of both models, we found no significant evidence to reject the NearlyNeutral model (p-value= 0.98416), suggesting that sequences in these sub-clusters of the AcrIF7 family are not under positive selection pressure. Furthermore, the Naive Empirical Bayes (NEB) results from the M1a model (Supplementary Data - M1a), which calculates the omega (dN/dS) values and their probabilities, showed that positions 29, 30, 42, 43, 45 and 64 (using *g2* as reference) have an omega of around 1, predicting that they have a neutral effect on the protein despite displaying sequence variation (Figure 3).

### Absence of a CRISPR-Cas system does not promote rapid accumulation of mutations in the AcrIF7 anti-CRISPR locus

The strains in which we identified AcrIF7 were very diverse in terms of ST, source and geographic location; besides, their isolation spanned over six decades (Figure 2). We wondered whether sequence conservation observed in AcrIF7 could be influenced by selective pressure imposed by CRISPR-Cas systems existing in the bacterial genome. Our search and classification of CRISPR-Cas systems in the *P. aeruginosa* genomes carrying *acrIF7* showed that around 40% of the isolates have at least one identifiable CRISPR-Cas system (47 out of 117 strains) (Figure 2F); a proportion similar to what has been reported previously [8]. From these, approximately 60% were subtype I-F (28 out of 47 encoding a CRISPR-Cas system). It is worth noting that in isolates with CRISPR-Cas I-F, only AcrIF7 of the subclusters sc1 and sc2 were identified (Supplementary Table S1).

We then investigated the effect of CRISPR-Cas presence on anti-CRISPR variation in an evolution experiment. We serially propagated the phage H70 (that carries *g2*) in either PA14 WT or PA14 ΔCR and determined the efficiency of the evolved phage lineages to evade the CRISPR-Cas system (Figure 5A). No statistically significant differences were found amongst any of the evolved stocks (p-value of 0.6468 for the phage evolved in PA14 WT and 0.7088 for the one propagated on PA14 ΔCR), indicating that the efficiency of AcrIF7 to inhibit the CRISPR-Cas system remained stable throughout the passages regardless of the absence of an active CRISPR-Cas system (Figure 5A).

**Figure 5.**
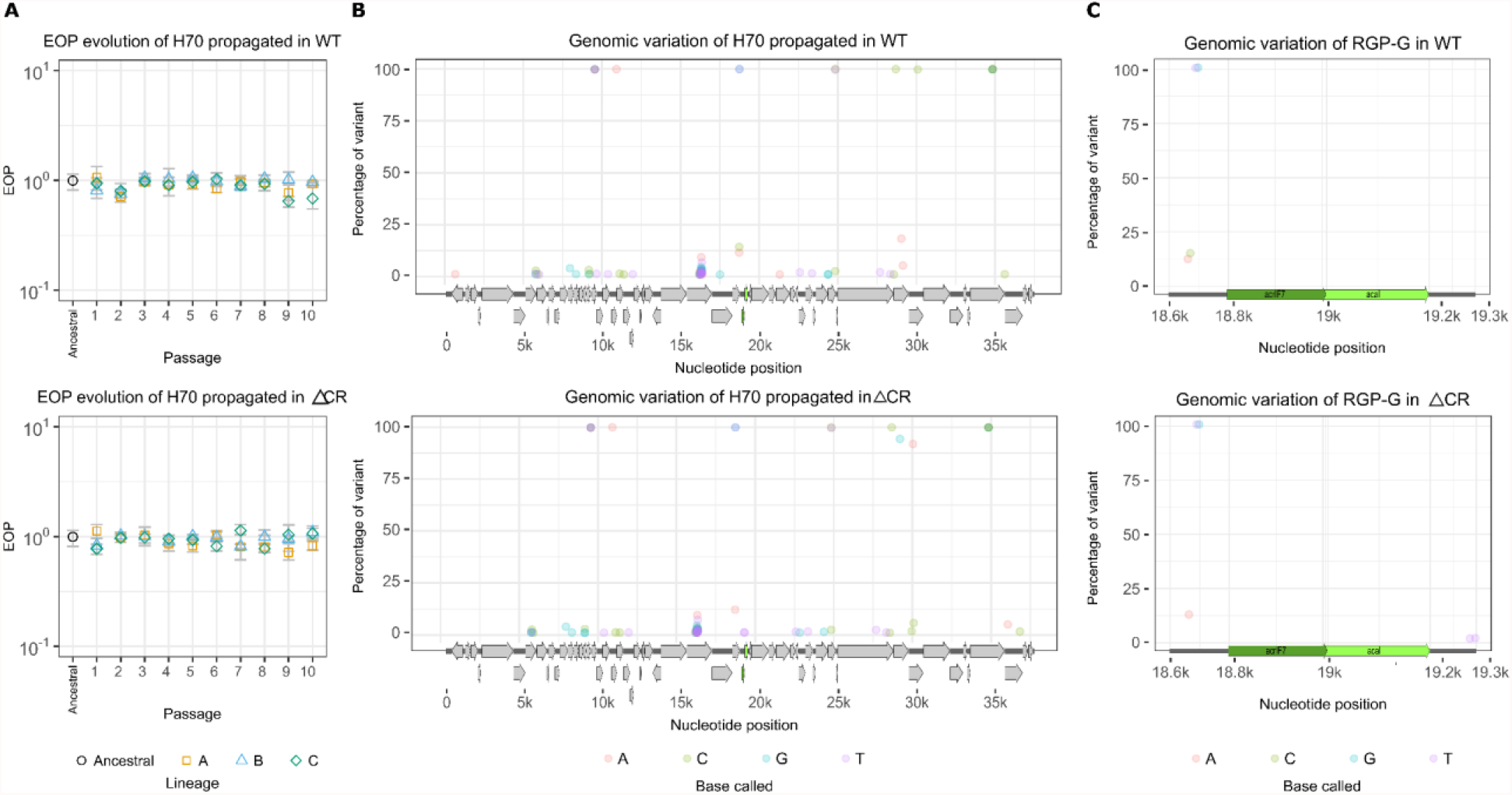
Experimental Evolution of H70 in PA14 WT and PA14 ΔCR. Panel A) illustrates the evolution of the efficiency of plating throughout the passages in either PA14 WT (top) or PA14 ΔCR (bottom). Each coloured shape represents a different lineage (biological replicates) with 3 technical replicates each. No statistically significant difference was found in one-way ANOVA tests. Panel B) shows all the variants in the H70 genome (from the last passage) that differ from the reference (NC_027384.1) and comprise more than 1% of the population. Panel C) represents the variants found in the region of genomic plasticity (RGP) G that is composed of the genes *acrIF7* (*g2*) and *aca1* (*g9*), and their respective intergenic regions.

Deep whole-genome sequencing and variant calling identified a few fixed mutations arising in the evolved H70 genome (Figure 5B and C): 15 in the phage propagated on PA14 WT and 16 in the phage evolved on PA14 ΔCR (Figure 5, Supplementary Table S2). Additionally, we observed emerging variants (<25% of the phage population) across the genome of both evolved populations. In total 78 and 74 mutations present in more than 1% of the population were detected in H70 propagated in the wild-type strain and in the ΔCR mutant (Supplementary Table S2), respectively. The most variable gene in both conditions was the ORF 31 encoding the portal protein, carrying an average of 35 variants (Supplementary Table S2). By contrast, *g2* was one of the most conserved genes in the phage, displaying no mutations in neither of the populations (Figure 5C). In the rest of the RGP-G, which comprises the genes *acrIF7* and *aca1*, we found 4 mutations in the population passaged on PA14 WT and 5 in the phage stock coming from PA14 ΔCR, all of them in either the upstream region of *acrIF7* or downstream *aca1* (Figure 5C and Supplementary Table S2). These mutations are not likely to impact the anti-CRISPR activity as they were not located in the coding region or nearby the promoter, thus correlating with the phenotypes from the EOP experiments. These results show that in the absence of the CRISPR-Cas system, AcrIF7 remains conserved for at least the duration of our evolution experiment.

### Mutational scanning of AcrIF7 revealed amino acids contributing to mutational tolerance and protein stability

The most abundant sub-clusters in the AcrIF7 family were highly similar and predicted to be under nearly neutral evolution. Moreover, no variants emerged in the AcrIF7 coding region during our evolution experiment. We, therefore, wondered whether the conservation of this family was due to the negative effect that mutations could have on the anti-CRISPR function. To explore this hypothesis, we undertook an unbiased approach similar to directed evolution to generate sequence diversity. We used a random mutagenesis strategy, in which we implement error-prone conditions during the PCR to promote the misincorporation of nucleotides by the polymerase (32), to explore the sequence landscape. Three different mutagenic conditions were applied to widen the error rate spectrum (Supplementary Table S3), which ranged from 0.6 to 1.4% per position. We then functionally characterised the resulting library detecting mutations that impaired the protein function and some others with neutral effect (Figure 6, Supplementary Figure S2 and S4). Finally, to understand the molecular basis of the loss of anti-CRISPR function, we performed docking analyses of the mutants in interaction with Cas8f.

**Figure 6.**
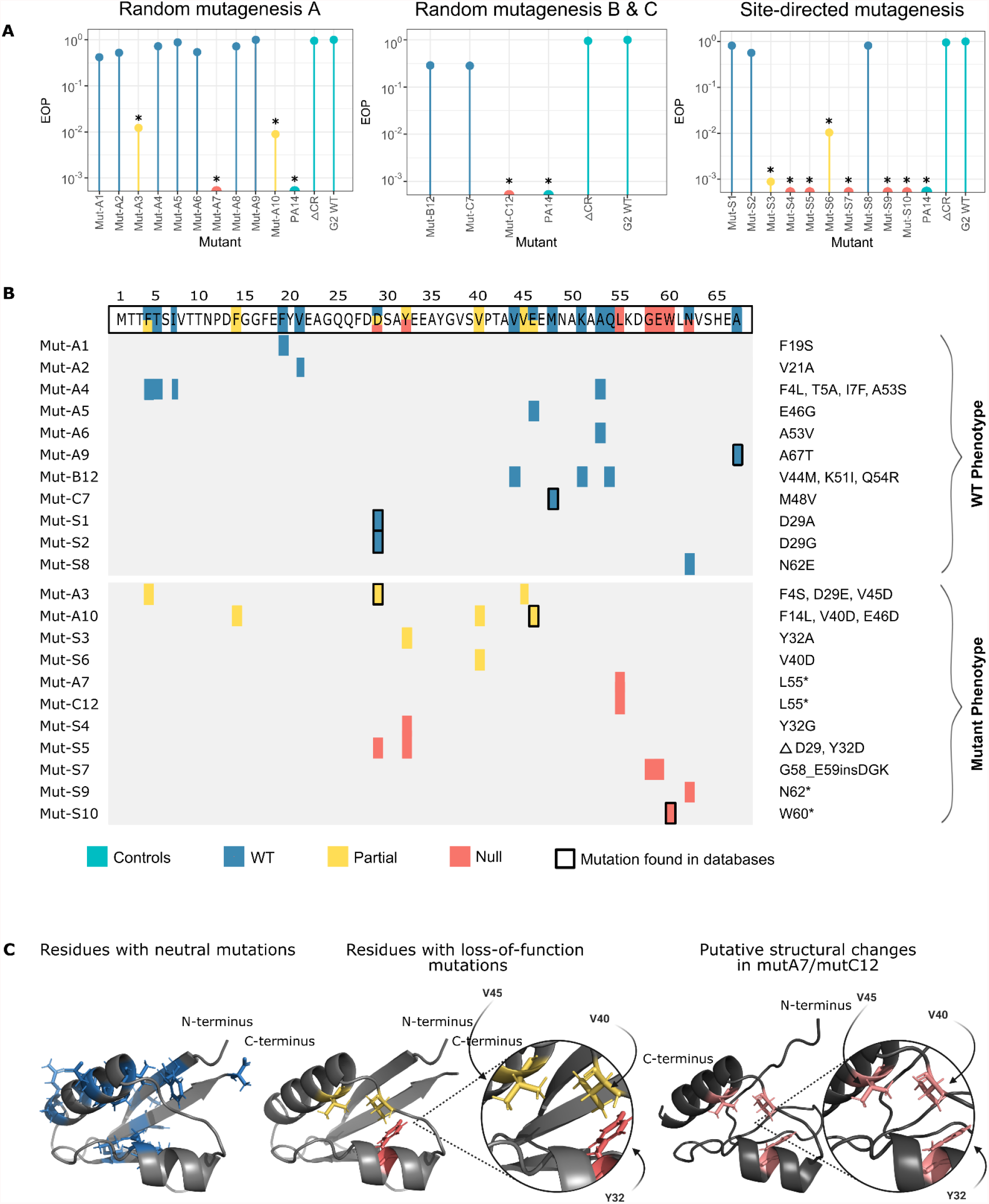
Impact of genetic variation on AcrIF7 function and structure. A) Efficiency of G2 mutants at inhibiting the CRISPR-Cas system I-F. The lollipop charts show the EOP of the CRISPR-sensitive phage JBD18 on PA14 carrying different variants of G2, normalised to the titre of the same phage in PA14 harbouring G2 WT. Asterisks denote adjusted p-values ≤0.05 (Raw data of replicates and p-values can be found in Supplementary Table S5). B) Mutational map of generated G2 variants. The colours represent the phenotype: wild-type (in blue), partial loss-of-function (in yellow), or null activity (in red). The changes in each mutant are shown next to the map (e.g. Mut-A1 has a mutation in F19S, whereas Mut-A3 has mutations in F4S, D29E, and V45D). The wild-type sequence of G2 is displayed at the top of the panel, with each mutated position coloured according to the phenotype of the mutant that carried changes in that position. Rectangles with a black bold contour indicate that those specific mutations (both position and amino acid change) were found in sequences in the databases. C) AlphaFold2 prediction of G2 structure showing residues with neutral mutations (in blue) or loss-of-function mutations (in yellow or red). Protein model prediction for the mutants mutA/mutC12 lacking 13 amino acids in the C-terminus. Amino acids in pink correspond to loss-of-function mutations; the figure shows the displacement of Y32 in the structure of the mutant, while V45 and V40 remain in the same predicted position as G2 WT.

Most of the tested mutant candidates displayed a wild-type phenotype (i.e., JBD18 was able to infect the strain carrying the mutant with the same efficiency as the strain containing the wild-type G2 - Supplementary Table S5), which harboured mutations scattered across the G2 sequence (Figure 6). Two mutants displayed a partial loss-of-function (Mut-A3 and Mut-A10). Although JBD18 EOP decreased ~100-fold in the partially functional mutants, the phage could infect the PA14 strain, indicating a residual protection effect provided by these G2 variants. Mut-A3 featured three punctual mutations: F4S, D29E, and V45D. The change in residue 29 of aspartate for glutamate is not expected to impact the protein function as both amino acids are chemically similar. As for the mutation F4S, a mutant with wild-type phenotype (Mut-A4) also featured a mutation in the same position (Figure 6), thus pinpointing the change in the valine 45 for an aspartic acid (V45D) as the most likely driver of the reduction in anti-CRISPR activity observed in Mut-A3. Similarly, Mut-A10 carried the mutations F14L, V40D, and E46D, from which the mutation V40D is expected to have a larger impact since a non-polar amino acid was replaced by an acidic one. Two mutants featured null anti-CRISPR activity: Mut-A7 and Mut-C12 (Figure 6). Both null mutants acquired mutations that resulted in the introduction of a premature stop codon in position 55 (L55*), albeit in a different manner (Figure 6). The premature stop codon resulted in the deletion of 13 amino acids of G2, which could potentially alter its tertiary structure (Supplementary Figure S6) and, therefore, the interaction with Cas8f.

Analysis of the structure model of G2 predicted by AlphaFold2 [40] showed that residues with identified neutral mutations, i.e., those that did not affect the anti-CRISPR activity, were dispersed in the protein structure (Figure 6C, residues in blue). On the other hand, residues with mutations that we predicted to be responsible for the partial loss of function observed in Mut-A3 and Mut-A10, namely V40 and V45, were located closely in the structure model (Figure 6, residues highlighted in red). Intriguingly, V40 and V45 were tightly clustered with a tyrosine located in the short alpha-helix (Y32) in the interior of the protein (Figure 6C). Therefore, we hypothesised this amino acid could also be necessary for the anti-CRISPR function. To test our hypotheses, we created a new series of mutants by site-directed mutagenesis (identified as Mut-S in Figure 6). We changed Y32 for A and G (Mut-S3 and Mut-S4, respectively) observing a 1000-fold reduction in the anti-CRISPR activity or no activity at all, respectively (Figure 6). According to our docking analyses, these two mutants are predicted to lose the interaction with R259 in Cas8f, which is part of the positively charged channel responsible of the non-target DNA displacement [51] (Figure 7). These results indicate that mutations in Y32 have a negative effect on the protein function and therefore explain why this residue is conserved among the members of the family.

**Figure 7.**
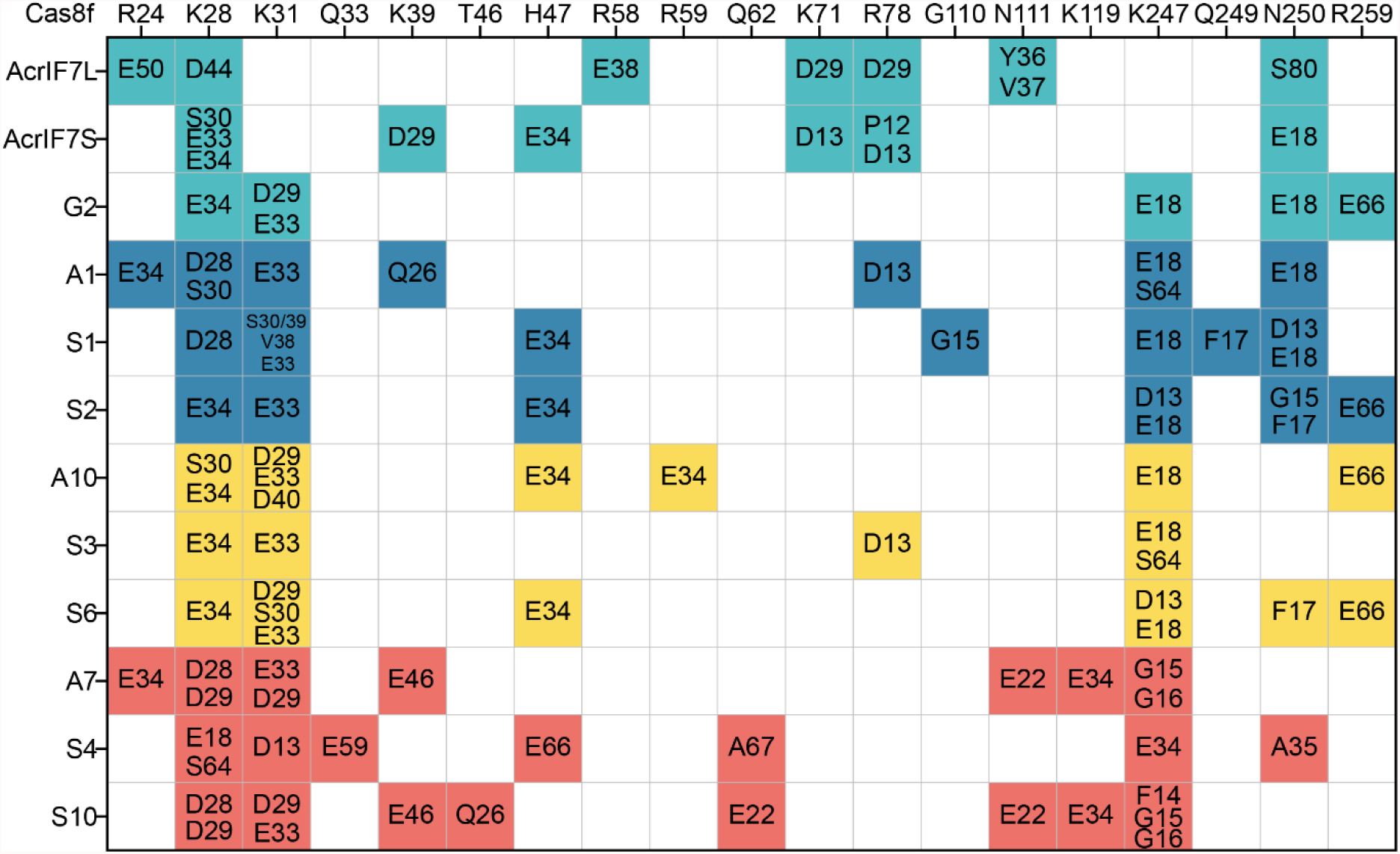
Residue-residue interactions of G2 and its mutants with Csy complex. Residue-residue interactions between the mutants and Cas8f. AcrIF7L [50], AcrIF7S [18] and G2 are coloured in green, mutants with wild-type phenotype are coloured in blue, mutants with partial loss of anti-CRISPR activity are coloured in yellow and mutants with null activity are coloured in red.

Next, we assessed whether the mutation V40D drove the partial loss of function previously observed in the mutant Mut-A10 (F14L, V40D, E46D), which carried two additional mutations. The new mutant Mut-S6 (V40D) exhibited a 100-fold reduction of the EOP (Figure 6). Similarly, the residue-residue interaction analyses revealed that that both MutS6 and Mut-A10 lost the AcrIF7:E18-Cas8f:N250 interaction, confirming that the residue V40 is responsible for the impairment of the anti-CRISPR activity. This is consistent with the observations of Guo *et al*., [52] who found that Cas8f N111 and N250 are responsible for the recognition of the PAM duplex on target DNA, and essential for CRISPR activity.

As one of the residues predicted to have a neutral effect on the protein, and identified as one of the most variable positions, was D29, we replaced it for A or G (Mut-S1 and Mut-S2, respectively). We observed no significant differences in the anti-CRISPR activity of these mutants, thus confirming that mutations in D29 do not influence the function (Figure 6). Moreover, it implies that the mutation V45D caused the partial loss of function in Mut-A3, likely due to missing interactions with N250 and R259 of Cas8f (Figure 7). Altogether, the evidence shows that Y32, V40 and V45 are functionally important. Together, they likely play an important role in stabilising the protein structure.

Our residue-residue interaction analyses of the mutants demonstrated that the interactions with K28, K31, N250 and R259 of Cas8f are conserved across the mutants. This suggests that the negatively charged surface of AcrIF7 is important for binding the positively charged channel of Cas8f and thus for anti-CRISPR activity, consistent with Kim *et al*. and Gabel *et al*. findings (Figure 7) [18, 50]. Our data shows that the loss of AcrIF7:E34-Cas8f:K28, AcrIF7:E18-Cas8f:N250 and AcrIF7:E66-Cas8f:R259 can lead to a reduced or abolished anti-CRISPR activity in loss-of-function mutants.

We further investigated the importance of protein length since this feature was variable amongst the members of the AcrIF7 family (Figure 3), and our null mutants Mut-A7/C12 were shorter in the carboxy-terminus (Figure 6). We introduced stop codons in the positions 60 and 62, and tested whether the variants were still functional. We found that mutants Mut-S9 and Mut-S10 that lost of 8 and 6 amino acids, respectively, were inactive (Figure 6). Interestingly, changing the residue in position 62 (N62E, mutant Mut-S8) had a neutral effect on the anti-CRISPR activity, implicating that loss of residues in the carboxyl terminus and the subsequent loss of a beta strand, which could destabilize the structure (Figure 6, S6), is the main factor driving the protein inactivation. Together with the analysis of the diversity of the AcrIF7 family, these results suggest that 67 amino acids constitute the minimal functional AcrIF7.

In summary, we introduced 30 different mutations in 21 positions scattered throughout the anti-CRISPR gene (Figure 6), corresponding to 31.3% of the protein. Seven of the mutations generated *in vitro* were present in AcrIF7 variants identified in databases (Figure 6B), with six of them displaying a wild-type phenotype. Notably, mutations introduced in 14 different positions, corresponding to 66.6% of the mutated residues, showed a neutral effect on the protein function, suggesting that these positions contribute to the mutational tolerance of AcrIF7.

## DISCUSSION

CRISPR-Cas systems are widespread in both archaea and bacteria [6], leading to the expectation that a wide variety of anti-CRISPRs exist in phage or other mobile elements to aid them in circumventing this defence system. Consistent with this notion, numerous anti-CRISPRs have been identified recently [12], and it can only be expected that the number of anti-CRISPRs will keep rising rapidly. Still, many questions remain unanswered regarding how these remarkable proteins work, evolve and spread. In this study we used AcrIF7 as a model to investigate diversity, distribution, evolution and functionality within an anti-CRISPR family.

By comparing homologs of the anti-CRISPR *g2* identified in different databases, we portrayed a detailed picture of the diversity within the AcrIF7 family. This enabled the identification of different sub-clusters and characterisation of their levels of sequence similarity, as well as distinguishing prevalent types and variants representing the diversity of the group (Figures 2 and 3). Our findings indicate that AcrIF7 homologs are mainly associated with *P. aeruginosa*, possibly suggesting specialisation to the CRISPR-Cas system of the species in which this anti-CRISPR family was first reported [45]. One exception is the member of the sub-cluster 3, a hybrid of the anti-CRISPR families IE4 and IF7 found in *P. citronellolis*, indicative of the potential flexibility of these molecules [46, 47].

Further exploration of regions flanking AcrIF7 homologs in complete genomes uncovered that this anti-CRISPR type can be linked to diverse genetic backgrounds in *P. aeruginosa* (Figure 4), indicating gene mobilisation. For example, we identified AcrIF7 homologs of the sub-cluster 1 in three unrelated phages sharing sequence similarity in less than 2% of their genomes: the siphophage H70 isolated from a clinical strain of *P. aeruginosa* in Mexico [21], the podophage Ab31 isolated from wastewater in Ivory Coast [53], and the siphophage TC7 isolated from hospital sewage in China (see metadata in GenBank record MG707188.1). The isolation source of these phages and the *P. aeruginosa* isolates carrying AcrIF7 (Figure 2) outlines the importance of anti-CRISPRs in different environments.

Although AcrIF7 homologs of the sub-cluster 1 were identified in distinct phages, they were predominantly associated with transposable phages of the type D3112-like (Figure 4). The genomes of these phages are known to contain multiple regions of genomic plasticity accommodating various accessory genes, with one region in particular harbouring diverse arrays of anti-CRISPR genes [10, 21]. In fact, anti-CRISPRs were first discovered in D3112-like phages [10]. It hence appears that this type of transposable phages represents a major reservoir of anti-CRISPR genes in *P. aeruginosa*, which could then be transferred to other mobile elements, consistent with our observation on the abundance of AcrIF7 homologs in D3112-like phages compared to other phage types. AcrIF7 homologs were also detected in divergent regions of *P. aeruginosa* genomes sharing little or no similarity with phage sequences, implying AcrIF7 acquisition by yet uncharacterised genetic elements. Unlike other anti-CRISPRs [13, 54, 55], we did not detect AcrIF7 homologs in virulent phages nor in plasmids. Whilst this observation may hint a preferred association with temperate phages or other types of mobile elements such as Integrative and Conjugative Elements, it could also be explained by a limited number of records existing for these elements in *P. aeruginosa*, especially for plasmids.

The most divergent sequences in the group AcrIF7 are closely related proteins encoded in *Janthinobacterium sp*. genomes, here clustered within the sub-cluster 5 (sc5; Figure 2). These sequences, however, are nearly identical to a protein that did not exhibit anti-CRISPR activity when tested against the CRISPR-Cas systems I-F and I-E of *P. aeruginosa* and I-F of *P. atrosepticum* [45]. This suggests that members of the sub-cluster 5 may either be active against CRISPR-Cas systems of other species or feature an unrelated function. The fact that proteins in sc5 share 62% similarity with those of the sub-cluster 1 (Supplementary Table S6), and can therefore be easily detected as potential homologs through BLAST searches, prompts us to be cautious about how we infer anti-CRISPR functions from sequence homology information. Since only representatives of the sub-clusters 1 and 3 in the group AcrIF7 have been experimentally verified as anti-CRISPRs ([14] (http://cefg.uestc.cn/anti-CRISPRdb/), it remains to be seen whether proteins in the sub-clusters 2 and 4 share the same function. In line with this remark, no Aca1 homologs were identified adjacent to representatives of sc2 and sc4 analysed here, hinting at either lack of anti-CRISPR activity for proteins in these subclusters or the presence of alternative Aca within the AcrIF7 family.

Random mutagenesis coupled with a selection method is a powerful approach to characterise proteins for which limited information is available [56]. This strategy is particularly effective at identifying not only residues involved in protein binding, but also in protein folding, or at finding mutations that enhance protein activity [36, 57, 58]. This is relevant for protein families such as AcrIF7, which are highly conserved according to the sequences available in databases, making it difficult to draw inferences about the impact that certain residues have on the protein. By implementing this approach, we not only captured and experimentally characterised some of the AcrIF7 diversity observed in databases, but generated more variation to study the functionality of a conserved anti-CRISPR. Remarkably, our mutational screening revealed important residues (Y32, V40, V45) contributing to the anti-CRISPR functionality that could not be identified with traditional methods which are focused on testing polar amino acids [18, 19]. Additionally, this approach enabled the identification of regions that contribute to the mutational robustness of the protein (Figure 6B, residues in blue).

Essential proteins are typically conserved [59]. This principle may help to understand why some anti-CRISPR families feature high levels of sequence similarity. Although not all phages encode an anti-CRISPR, this function becomes essential when infecting a host with a CRISPR-Cas system. In this context, mutational robustness represents an advantageous trait that minimises the effect of random changes on the protein function [60, 61]. Despite the high levels of similarity observed in most of the members of the AcrIF7 family, our experiments identified some mutations with a neutral effect on the anti-CRISPR function. These residues contribute to the mutational tolerance of the protein, corresponding to ~67% of the amino acids mutated in our study. Our findings are reminiscent of those reported for the Influenza A virus matrix protein M1, and the middle domain of the heat shock yeast protein Hsp90 [62, 63], for which high levels of intrinsic tolerance to mutations were found despite the little sequence variation observed in nature. These proteins play crucial roles in the survival of the microorganism carrying them: AcrIF7 is essential for a successful phage infection in the presence of a CRISPR-Cas system; M1 participates in multiple stages of the viral infectious cycle; Hsp90 is involved in protecting cells from environmental stress and growth at high temperature [62, 63]. Future studies focused on deeper mutational scanning of anti-CRISPRs wil tell us how robust they are in comparison with other proteins.

Although G2 was capable of carrying multiple mutations without having its function significantly affected (Figure 6, Supplementary Table S5), it is possible that those variants are not as stable as the wild-type version and are therefore negatively selected; thus suggesting that the G2 sequence represents the optimal AcrIF7 version. This notion is supported by the fact that G2 can completely block the CRISPR-Cas system I-F, i.e., a CRISPR-sensitive phage can infect a strain carrying G2 WT with the same efficiency as the mutant lacking the CRISPR-Cas system (Figure 6b). Moreover, variants sharing the G2 sequence are the most prevalent among the members of the family (Figure 3), possibly suggesting a selection for this version of AcrIF7.

Studies by Kim *et al*. and Gabel *et al*. found that AcrIF7 binds the Cas8f–Cas5f complex and that the interaction was driven by hydrogen bonds and electrostatic interactions [18, 50]. Mutations in D28, D29 and D57 have been reported to affect the binding affinity by 3 to 7-fold while D13, E33 and E34 by more than 60-fold [18]. It has been experimentally verified that D13 interacted with K71 and R78 in Cas8f, D28 interacted with K28 while E34 interacted with R24 [50]. However, the mutations in D29 in this study (D29A in Mut-S1 and D29G in Mut-S2) displayed wild-type phenotype (Figure 6), suggesting that AcrIF7 can maintain the phenotype with the conversion from negatively charged residue to hydrophobic residues in D29. This position is quite variable in nature where D29A exists at a 4-fold higher frequency than D29G (Figure 3). Likewise, the AcrIF7 variant carrying the mutation E46K binds Cas8f with similar affinity as the wild-type [18]. Similarly, no change in the phenotype was observed in our mutant harbouring the mutation E46G (Mut-A5) (Figure 6).

We discovered that Y32 is key for the anti-CRISPR activity of G2. Based on its location inside the short alpha-helix, and our residue-residue interaction analysis predicting no hydrogen bonds, we hypothesise that this residue contributes to the anti-CRISPR activity by stabilising the protein. Likewise, we uncovered that mutations in V40 and V45 had a significant impact on the function, likely because of their contribution to the protein hydrophobic packing [64]. This highlights the importance of mutational screening strategies to study anti-CRISPR proteins, as it allowed us to find amino acids that are not part of the protein surface but still contribute largely to its function. Our work also underscores the importance of functional analyses to complement structural studies, as we identified mutations in residues reported to interact with the CRISPR-Cas system (V21, D29) [18, 50] which had neutral effect on the anti-CRISPR function.

We observed that deletions in the Carboxy-terminus of G2 completely abolished the anti-CRISPR function. The structure model of the mutants with the deletion L55* (Mut-A7/C12) suggests that this region is important for the maintenance of the structure and the stability of the protein (Figure S6). We did not identify longer AcrIF7 variants on the Carboxy-terminus side in databases. In contrast, we found that the anti-CRISPR encoded in the genome GCF_900707915 is shorter. Our characterisation of the mutant Mut-S9 (N62*), however, suggest that this variant is no longer active. Together, our mutants characterisation and bioinformatics analyses indicating that the predominant variant length is 67 amino acids (93.6% of the sc1 and nearly 65% of all sub-clusters), suggest that this protein size has been evolutionarily selected.

Besides furthering our understanding of the phage-bacteria arms race and co-evolution, the study of anti-CRISPRs can boost their application in biotechnology, similar to the recent boom in the development of CRISPR-Cas-based technologies [65]. For example, it has been proposed that anti-CRISPRs can aid the use of CRISPR-Cas9 systems to control gene expression as regulatory tools preventing off-target editing [66]. Additionally, we envisage that anti-CRISPRs can be powerful tools in the fight to control multi-drug resistant bacteria by providing phages engineered for therapy purposes with a gene repertoire enabling them to evade the CRISPR-Cas defence of the target pathogen and increasing their host range. To this end, it is necessary to not only continuing the discovery of new anti-CRISPRs but to thoroughly characterise them and identify their optimal versions for biotechnological use. We consider that a strategy like the one presented in this study, i.e. combining genomics and phylogenomics analyses, mutational scanning and functional characterisation, is a first step towards that direction.

## DATA AVAILABILITY

All sequences analysed in this work are publicly available from anti-CRISPRdb or GenBank. The corresponding accessions are indicated throughout the text and in figures 2, 3 and 4. The sequences from the evolution experiment can be found under the BioProject ID PRJNA796330.

## ACKNOWLEDGEMENTS

We thank Guadalupe Aguilar González from Unidad de Ácidos Nucleicos, Department of Genetics and Molecular Biology, and Dr Dulce Delgadillo Álvarez from Unidad de Genómica, Proteómica y Metabolómica - CINVESTAV, for technical assistance in Sanger sequencing and Miguel Ángel Moreno Galeana and Dr Eva Jacinto from DGMB, CINVESTAV, for technical support with lab experiments. We also thank Dr Herminia Loza Tavera from the Faculty of Chemistry, UNAM, for providing us with pUCP24 plasmid, and Prof Alan Davidson for sharing with us the strains PA14 WT, PA14 ΔCR and the phage JBD18.

## AUTHORS CONTRIBUTION

W.F. and A.C. conceptualised the study and drafted the manuscript with input from other authors. W.F. carried out the positive selection analysis, experimental evolution, random and site-directed mutagenesis, functional experiments, Sanger and Illumina sequencing, and analysis of the protein structure. A.C. performed the homologs search, analysis of metadata, phylogenetics analysis, sequences comparison and comparative analysis of flanking regions. D.C. contributed to the conceptualisation of the study, performed infection assays, and helped draft the manuscript. Y.W. performed the structural analysis of G2 wild-type and mutants and contributed to drafting the manuscript. A.D.C. cloned and tested the function of G2 WT. M.W. helped design the evolution experiment and analyse the protein structure. G.G. and L.K. contributed by supervising the work and revising the manuscript. F.L.N. helped with the design of the experimental evolution, supervised the work carried out by Y.W., and contributed to drafting and revising the manuscript. G.G. was responsible for funding acquisition.

