## Supplementary Figures for "Distribution and molecular evolution of the anti-CRISPR family AcrIF7"

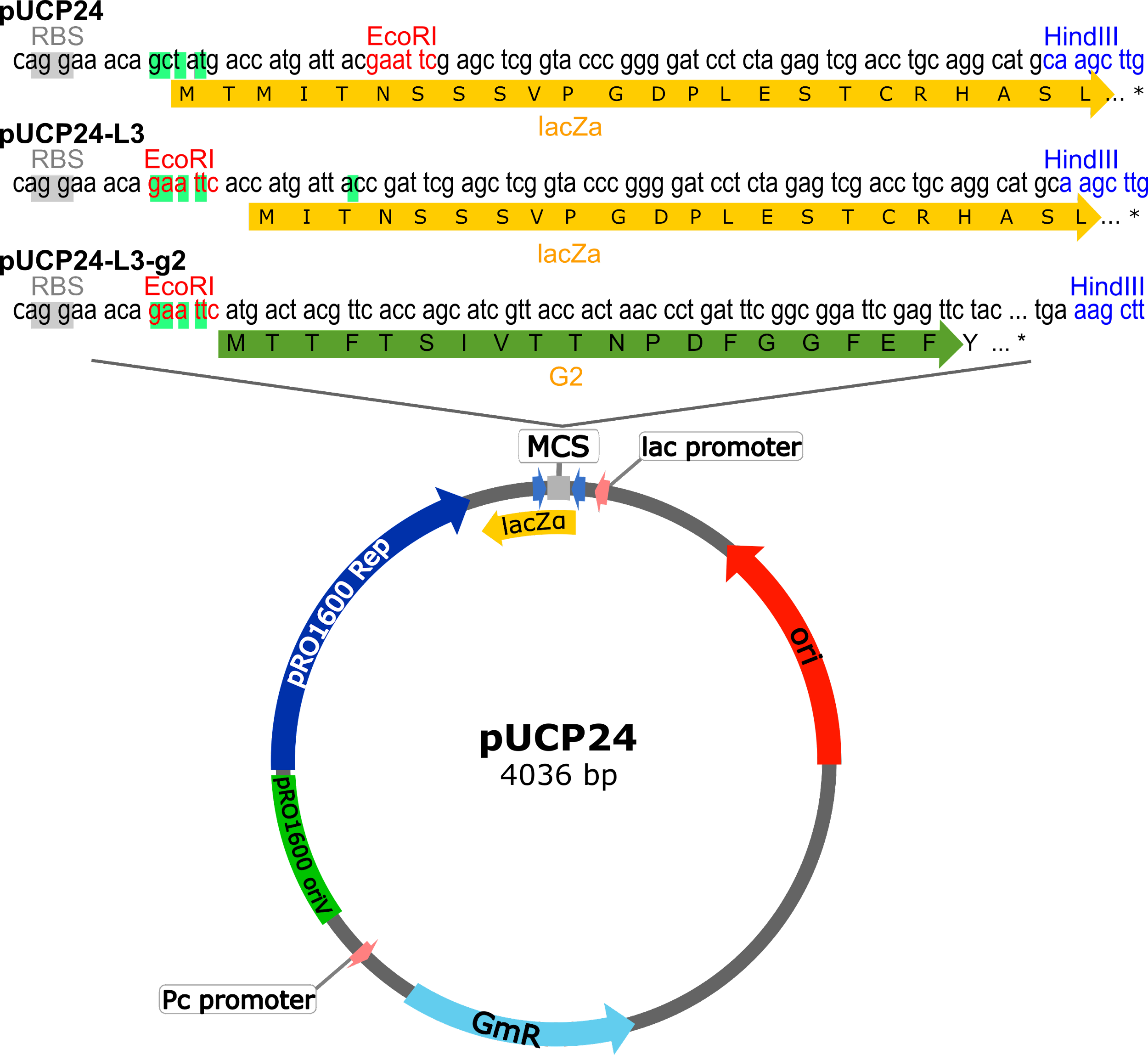

**Figure S1. Modified version of the expression vector pUCP24.** The vector map shows a modified version of the pUCP24 plasmid (named pCUP24-L3). The changes made to the original plasmid sequence are indicated in light green boxes. The yellow (lacZα) and green (*g2*) arrows represent the coding regions in the MCS of the plasmids. The changes in the plasmid involved moving the EcoRI restriction site in pUCP24 upstream to prevent the incorporation of additional amino acids from lacZα peptide into G2 once it is cloned.

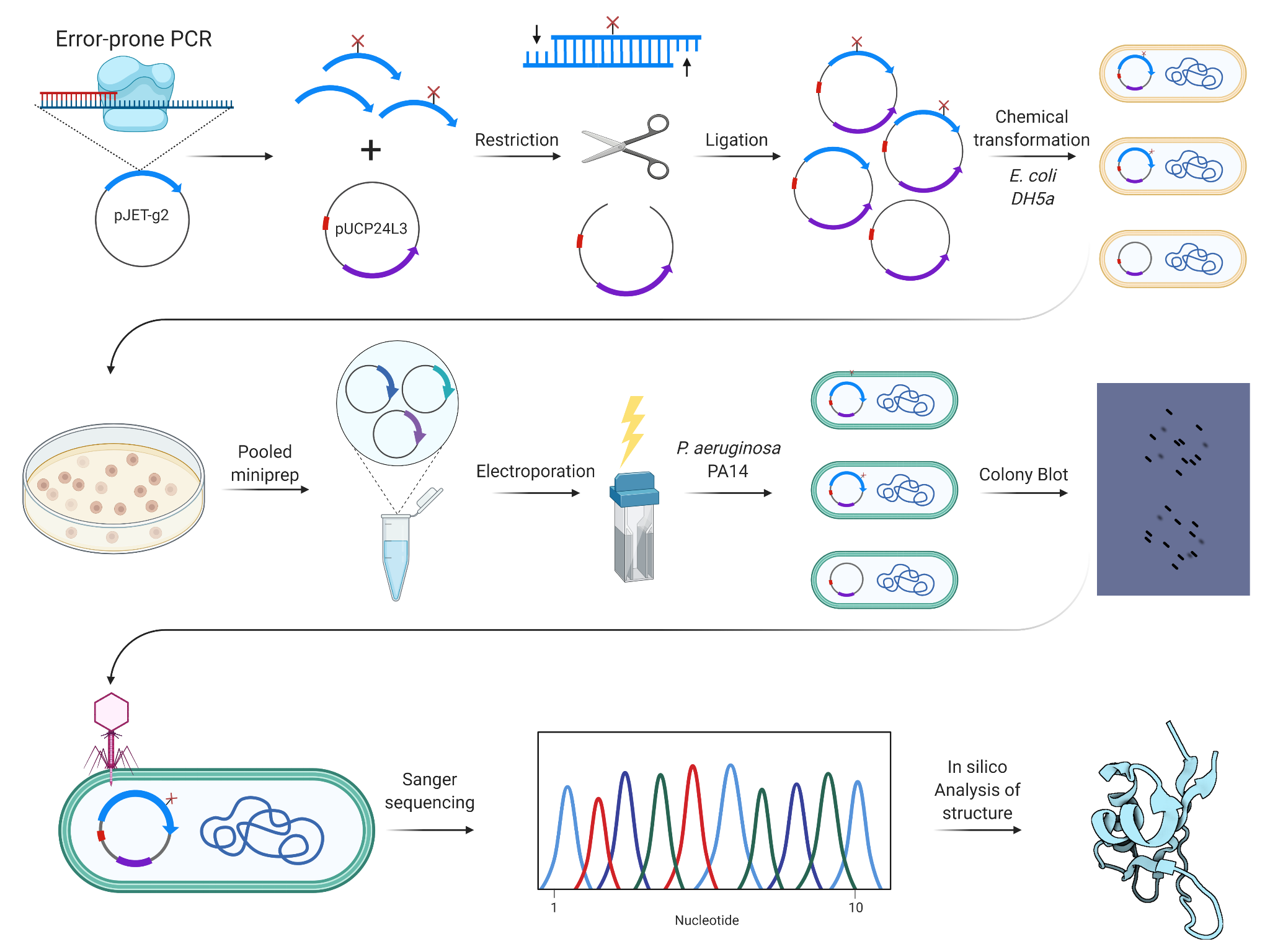

**Figure S2. Strategy for cloning and identification of G2 mutants.** The figure illustrates the steps followed to generate the collection of G2 random mutants presented in this study. The strategy consisted of 1) cloning the error-prone PCR products into pUCP24-L3, 2) transformation and extraction of pools of plasmids from *E. coli*, 3) electroporation of the pools into *Pseudomonas aeruginosa* PA14, 4) assessment of the efficiency of the variant to block the CRISPR-Cas system and 5) sequencing of the mutants and analysis of the protein model. Figure created with BioRender.com.

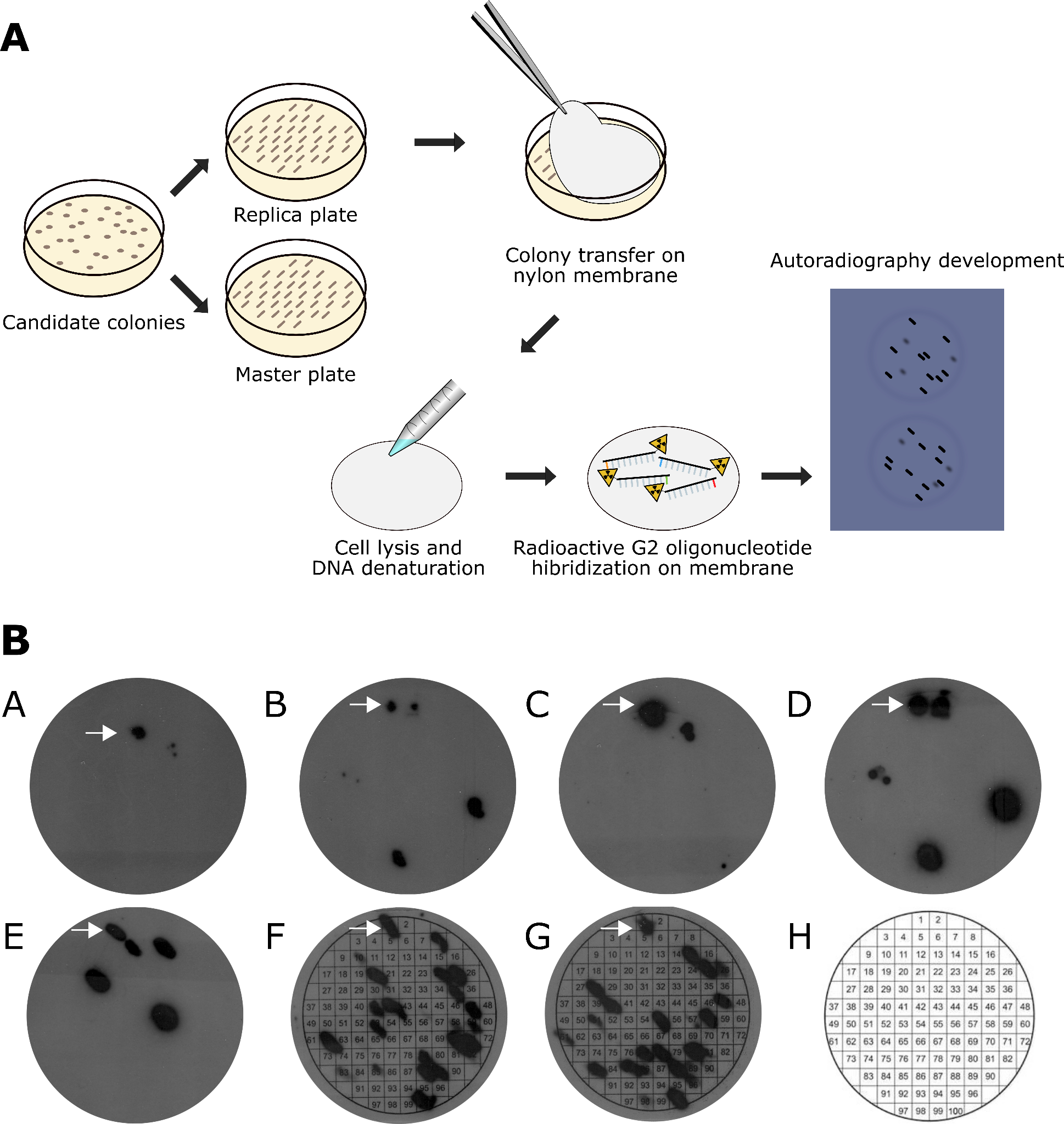

**Figure S3. Colony blot protocol.** I) The protocol consisted of streaking candidates in 2 LB plates (master and replica plates), followed by transferring the colonies to a nylon membrane. The membrane is processed with different solutions (see Methods) and hybridised with a radioactive probe. X-ray films are exposed to the membranes and developed to identify colonies with *g2*. II) The lower panel shows the x-ray films from 7 different membranes. Black spots confirm the presence of g2 in the colonies, which are then identified based on the position in the membrane (with the help of a grid template that was also used when streaking the colonies-H). The white arrows point to the positive controls included in each membrane.

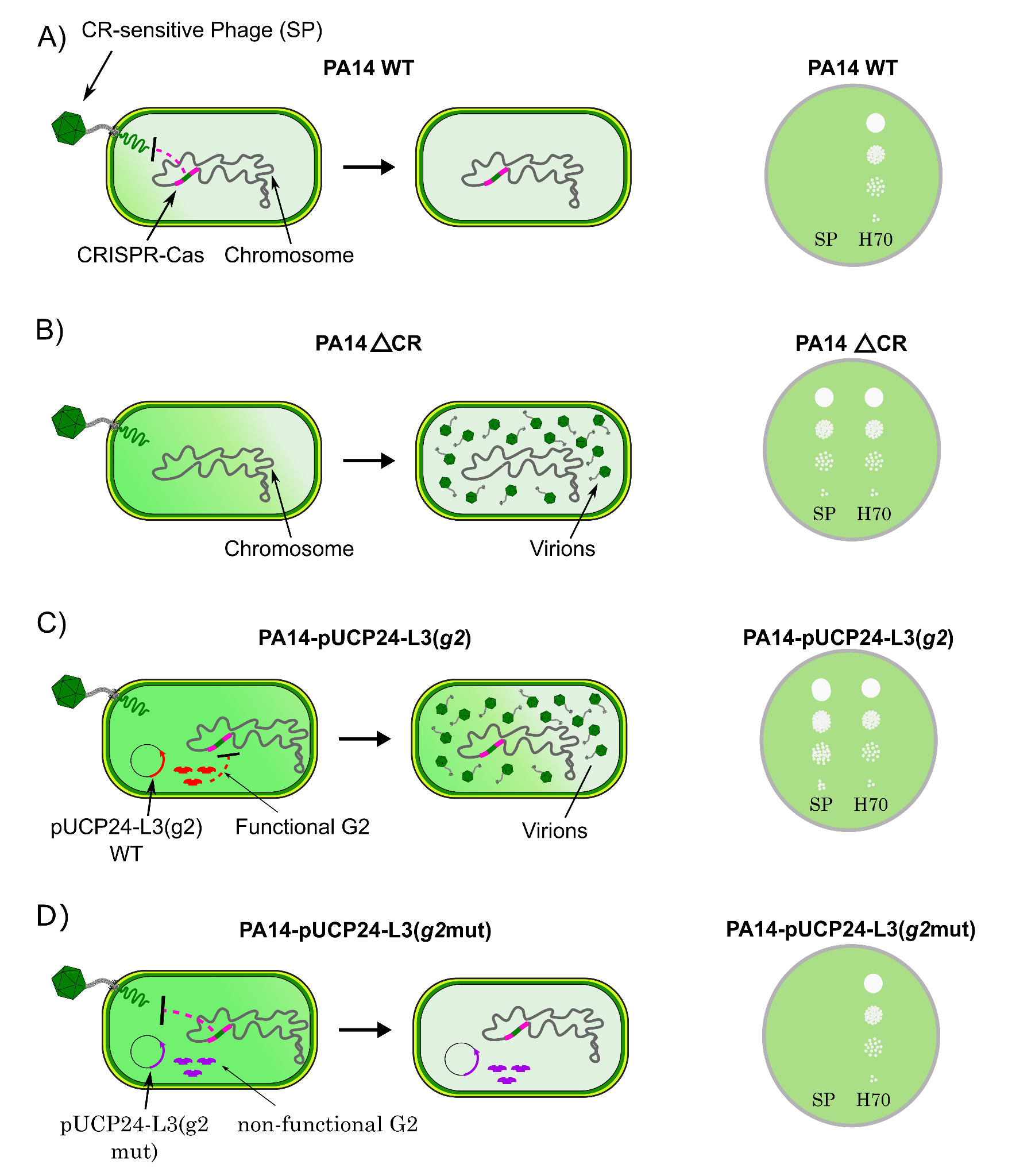

**Figure S4. G2-phenotyping assay.** The activity of G2 was assessed based on the ability of the variant to block the CRISPR-cas system and allow the infection of a CRISPR-sensitive phage. In figure 4 different scenarios are shown: A) PA14 with an active CRISPR-cas system that blocks the infection by a CRISPR-sensitive phage (SP), B) PA14 ΔCR mutant that can no longer defend against SP C) PA14 WT transformed with a wild-type version of the anti-CRISPR G2, which inhibits the CRISPR-cas system and therefore allows the infection the SP, and D) PA14 WT carrying mutant versions of G2 that are defective at suppressing the CRISPR-cas system, hence SP cannot infect the cell. The left panel shows the different scenarios at the cellular level and the right panel illustrates the phenotypes seen in bacterial lawns.

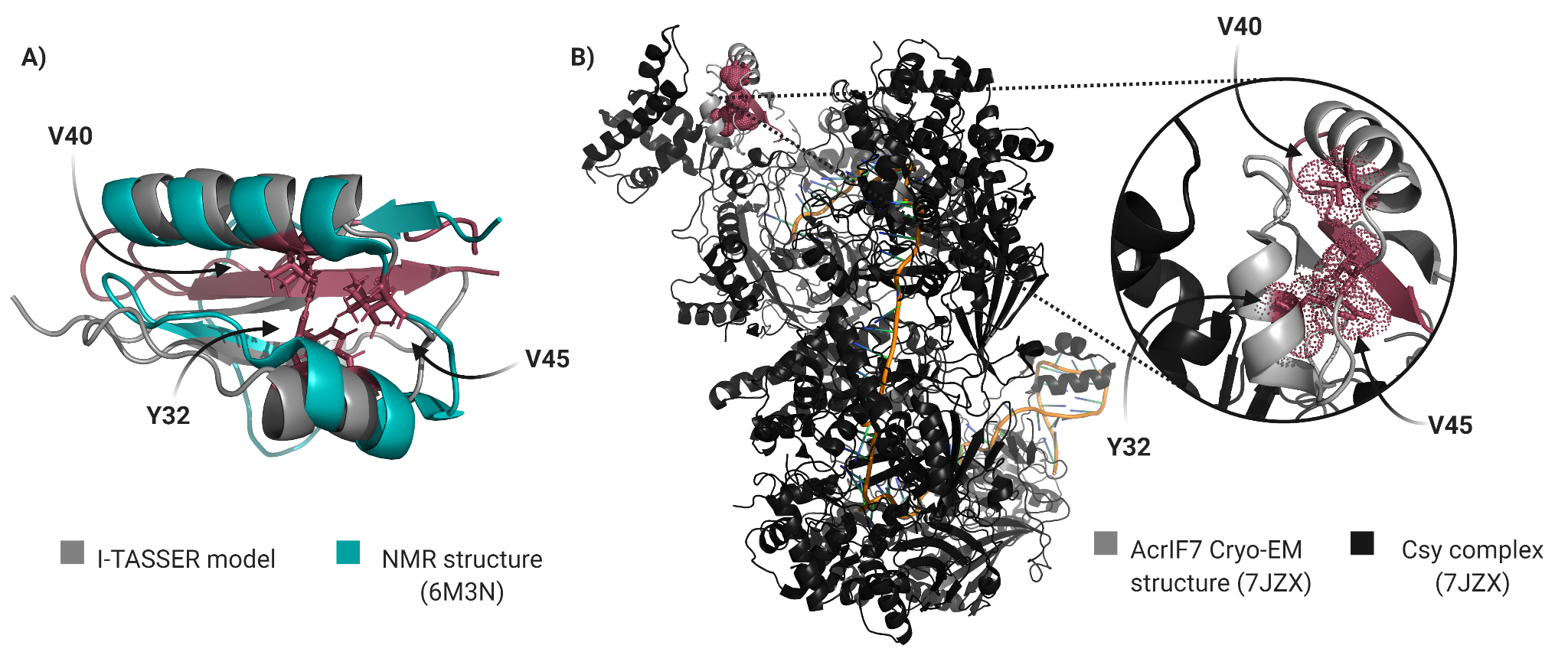

**Figure S5. Superposition of I-TASSER model and NMR structure.** A) I-TASSER protein model of G2 (in grey) and NMR structure (PDB 6m3n, in dark cyan) were overlaid using the “super” function in open-source Pymol, with an RMSD of 3.70 Å. Amino acids found to be important for anti-CRISPR activity are shown as sticks in burgundy. B) Cryo-EM structure of AcrIF7 in complex with Csy proteins (PDB 7JZX). The figure shows in burgundy the positions in AcrIF7 of the amino acids reported in the present study as important for the function.

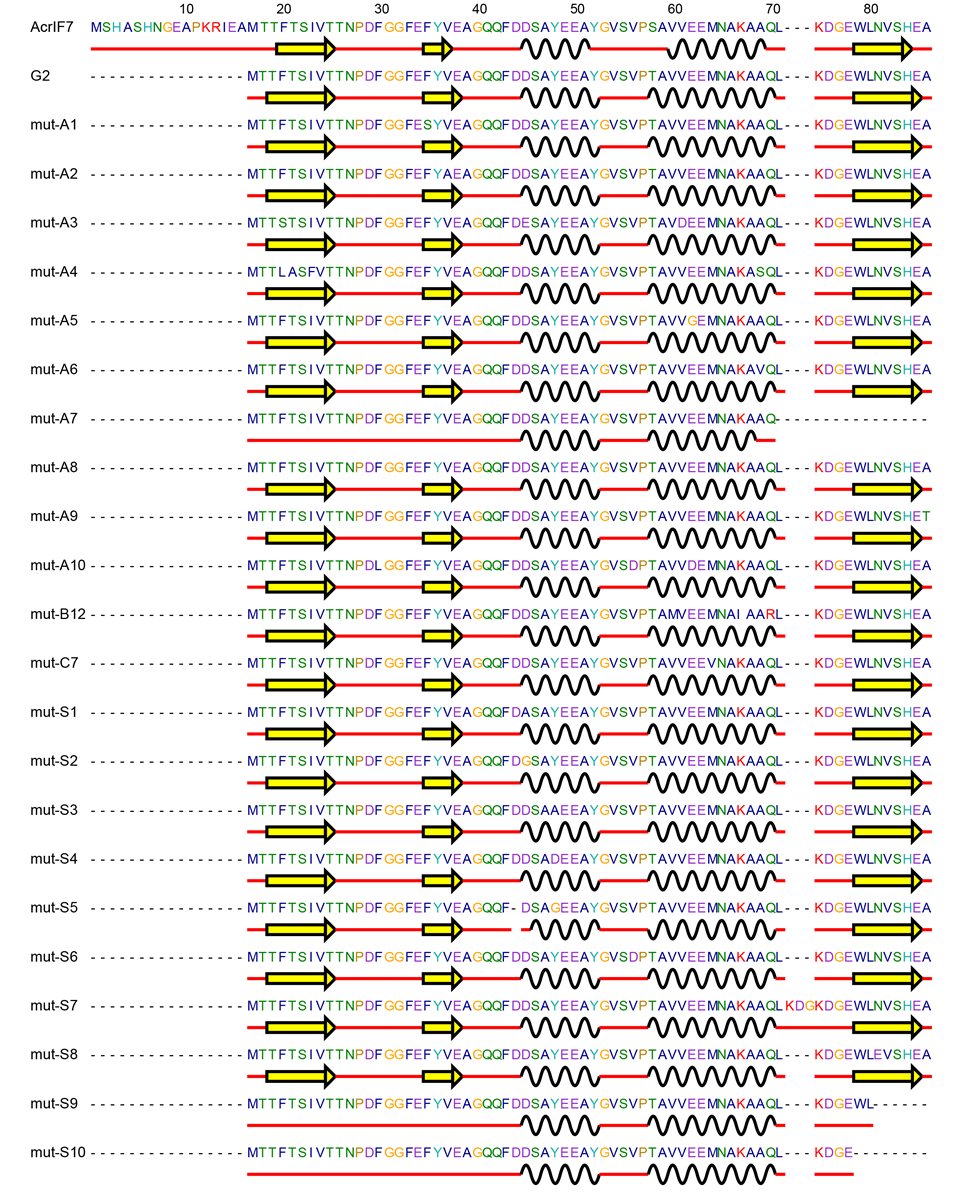

**Figure S6. The secondary structures of AcrIF7, G2 and its mutants.** The beta-sheets are represented as yellow arrows, while the alpha helices are represented as black wavy lines.
